# The multiomics landscape of small peptides encoded by long non-coding RNA-derived sORFs in rice

**DOI:** 10.1101/2025.04.01.646714

**Authors:** Zhipeng Chen, Haocheng Lin, Lirong Wei, Sunxi Wang, Ting Peng, Wei Song, Dacheng Wang, Yufeng Wu, Liuji Wu, Jingni Wu, Yiming Wang

**Author notes:** **Correspondence to** (Y. Wang); (J. Wu); and (L. Wu). The author(s) responsible for distribution of materials integral to the findings presented in this article in accordance with the policy described in the Instructions for Authors (https://academic.oup.com/plcell/pages/General-Instructions) is: Yiming Wang.

## Abstract

Long non-coding RNAs (lncRNAs), representing the non-coding RNA regions, constitute a significant portion of the genomes in complex organisms. Recent studies suggest that some lncRNAs have the capability to encode peptides. However, the presence of the lncRNA-derived sORFs-encoded polypeotides (LSEPs) in plants is not well understood. In this study, we developed a multi-omics approach that encompasses transcriptomics, translatomics (Ribo-seq), and proteomics to identify of LSEPs in rice. Among the 2764 identified lncRNAs, 42.69% were found to be bound by the ribosome, indicating a potential for encoding. Optimized small peptide extraction protocol was further developed, and the small peptides from rice leaves were extracted and subjected to LC-MS/MS analysis, leading to the identification of a total of 403 LSEPs across four constructed search databases. This work confirms the peptide-coding ability of lncRNAs in plants. Collectively, our study establishes an efficient multi-omics method for identifiying small peptides encoded by lncRNAs, which may be valuable for large-scare screening of LSEPs in plants.

## Introduction

Eukaryotic genomes produce a large number of non-coding transcripts during transcription (Mercer et al., 2009; Mattick et al., 2023). Non-coding RNAs, description as a class of RNA molecules transcribed from the genome but not undergoing translation, were previously believed to have minimal or no protein-coding capacity, despite being biologically functional (Mercer et al., 2009). Under the “one-gene one-polypeptide” paradigm, the canonical protein-coding sequence of mRNA is defined by the longest open reading frame (ORF) that includes an AUG start codon. Conversely, RNA transcripts exceeding 200 nucleotides without conserved ORFs are categorized as long noncoding RNAs (lncRNAs) (Mercer et al., 2009). lncRNAs have garnered significant interest due to their crucial roles in various cellular processes, including gene expression regulation, chromatin structure modification, epigenetic control, RNA processing and stability (Yap et al., 2010; Gong and Maquat, 2011; Luo et al., 2016; Isoda et al., 2017; Mattick et al., 2023). lncRNAs constitute a diverse class of functional ncRNAs. Linear lncRNAs are categorized based on their location related to nearby protein-coding genes into long intergenic ncRNAs (lincRNAs), intronic ncRNAs (incRNAs), and natural antisense transcripts (NATs) (Ma et al., 2013; Mattick and Rinn, 2015; Chen et al., 2017). Moreover, circular RNAs (circRNAs), which mostly originate from protein coding regions or intronic regions, represent another group of lncRNAs (Jeck and Sharpless, 2014). LncRNAs exhibit regulatory properties, with trans-acting lncRNAs operating at genomic locations distant from their transcription sites and performing regulatory functions elsewhere, while cis-acting lncRNAs influence gene expression within or near the same chromosomal region by targeting protein-coding genes(Yan et al., 2017; Gil and Ulitsky, 2020).

Emerging studies highlight that lncRNAs are key regulators in the fine-tuning of gene expression at the transcriptional or post-transcriptional level in almost all cellular processes (He et al., 2019; Statello et al., 2021). However, the roles of plant lncRNAs and their mechanisms of regulating cellular processes are not well understood. Although research is still in its early stage, the biological functions and mechanisms of a few plant lncRNAs have been reported as novel regulatory control switches in plant developmental (Zhou et al., 2021), flowering (Jin et al., 2023), abiotic stress responds (Zhao et al., 2023), and biotic tolerances (Liu et al., 2022; Zhang et al., 2022; Ai et al., 2023). The action modes of plant lncRNAs are diverse and complex. Similar to the reports in animal kingdom, plant lncRNAs can act as scaffolds and/or decoys to regulate transcription through chromatin modifications, by binding to DNA and/or protein (Heo and Sung, 2011; Engreitz et al., 2016; Roulé et al., 2022).

Benefiting from the development of next-generation sequencing technology, millions of sORFs imbedded in lncRNAs have been discovered. This raises the questions of whether those lncRNAs can function as mRNAs to translate functional peptides, and if so, how many proteins or peptides are encoded by these lncRNA sORFs. Obviously, the coding potential of lncRNAs has historically been overlooked. The length of sORFs-encoded polypeotides (SEPs) often fails to meet arbitrary cutoff, such as 100 amino acids, and is treated as “noise” or false-positives. Consequently, the roles of SEPs in cellular processes have been consistently ignored. Recently, several human lncRNA-transcribed SEPs have been validated and are proven to play important biological and pathological roles in muscle regeneration, immunity, inflammation, and tumorigenesis (Niu et al., 2020; Ge et al., 2021; Barczak et al., 2023).

To substantiate the coding capabilities of lncRNAs, researchers employ a multitude of methodologies, including Ribosome profiling (Ribo-seq), CRISPR-Cas9-based screens targeting sORFs/mutated sORFs, and LC-MS/MS proteomics. Ribo-seq is a deep sequencing-based approach that captures ribosome-protected mRNA fragments, offering a snapshot of translation. These ∼30-nucleotide ribosome footprints are converted into a library of DNA fragments and sequenced. Analysis of Ribo-seq reveals a 3-nucleotide periodicity indicative of translating ribosome’s progression along the mRNA, representing the stepwise movement of 80S ribosomes actively translating ORFs (Brar and Weissman, 2015; McGlincy and Ingolia, 2017). Coupled with sequence analysis, Ribo-seq yields a precise map of translated ORFs, facilitating *de novo* ORF discovery. Notably, Sonia et al. identified 7767 high-confidence sORFs, with CUG and GUG emerging as prevalent alternative start codons instead of AUG across various human cell types and tissues (Chothani et al., 2022). This start codon frequency distribution has also been confirmed by other approaches (Zhang et al., 2021). By integrating Ribo-seq with lncRNA sequencing, several lncRNAs with coding potential have been identified. Among them, lnc-AP, which produces a peptide (pep-AP) implicated in colorectal cancer cell apoptosis and sensitization to chemotherapy (Wang et al., 2022). Zhang et al. innovated this process by constructing a comprehensive lncRNA-derived SEPs (LSEPs) database and implementing a tailored enrichment strategy, leading to the discovery of 762 new LSEPs, marking a substantial contribution to the field (Zhang et al., 2021). Despite these advancements, the biological functionality of many predicted peptides, particularly those derived from plant lncRNAs, remains unexplored. To our knowledge, only a handful LSEPs, such as lncRNA7, have been functionally characterized (Zhang et al., 2022). To explore the LSEPs in plants, we hypothesize that the model plant *Oryza sativa* harbors a wealth of overlooked lncRNA-encoded ORFs with translational potential, awaiting exploration to unravel their biological significance.

In this study, we established a multi-omics workflow, consisting of transcriptome, translatome, and proteome, to identify the portion of potential translated rice lncRNA transcripts and to draft the rice SEPome for the first time. This approach enabled us to build a robust SEP database that includes all putative sORFs from lncRNA transcripts engaged in the active translation process, significantly improving the discovery of LSEPs in rice plant. A total of 403 OsLSEPs were identified using MaxQuant and Proteome Discoverer (PD) software from rice leaf proteins extracted by two different small peptide enrichment buffers. Our results demonstrate an experimental strategy that integrates small peptide extraction, LC-MS/MS identification, database establishment, and searching, which benefits the identification of small molecular weight peptides. Additionally, a preference for non-AUG start codons was also observed, as mentioned before. We then employed multiple technologies to experimentally validate the translation ability of these ORFs with different start codons (e.g. AUG, CUG and TUG), suggesting that a fraction of lncRNAs-sORFs is indeed translated in rice. These insights will not only enhance our understanding of the roles and actions of lncRNAs in plant, but also provide a multifaceted tool for exploring LSEP function in rice.

## Results

### The workflow for identifying lncRNA-encoded small peptides

To detect the small peptides encoded by lncRNAs in the whole rice genome, we developed an integrated analysis workflow combining transcriptome (RNA-seq), translatome (Ribo-seq), and proteome (LC-MS/MS) approaches (**Fig. 1A**). The expressed lncRNAs were first detected using next-generation sequencing from rice seedling leaves. On average, the Illumina sequencing platform yields 75 million reads, and 98% of these reads can be mapped to the rice reference genome (TIGR7) (**Supplementry Table 1A**). For successful encoding, the expressed lncRNAs must be bound by the ribosome. Ribosome-protected lncRNA fragments were detected by Ribo-seq, which revealed an average of 47 million reads per sample replicate (**Supplementry Table 1B)**. After filtering out housekeeping RNAs (rRNA, tRNA, snoRNA), 84% of the retained reads mapped to the reference genome, suggesting high-quality sequencing data. The size of ribosome-protected fragment (RF) is mainly between 28 and 31 nt (**Supplementary** Fig. 1A), similar to that previously reported for maize and animals(Lei et al., 2015; Yan et al., 2021). Importantly, the distribution of 31 nt RFs from rice exhibited a strong 3 nt periodicity when analyzed using the first base position of RFs, which is a typical feature of activate translation where the ribosome moved down 3 nt to recognize next codon (**Supplementry Fig. 1B**). Reading frame analysis of 31 nt RFs revealed that more than 50% of them accumulated in the first frame (**Supplementry Fig. 1C**). A high correlation was observed between the replicates of each sample in mRNA-seq libraries and Ribo-seq libraries (**Supplementry Fig. 1D**). Based on our results, 42.69% of the detected lncRNAs in rice leaves could be bound by the ribosome. Together, our data reveal a high-quality Ribo-seq RF features, which are consistent with previous reports in plants (Guo et al., 2023; Zhu et al., 2023).

**Figure 1.**
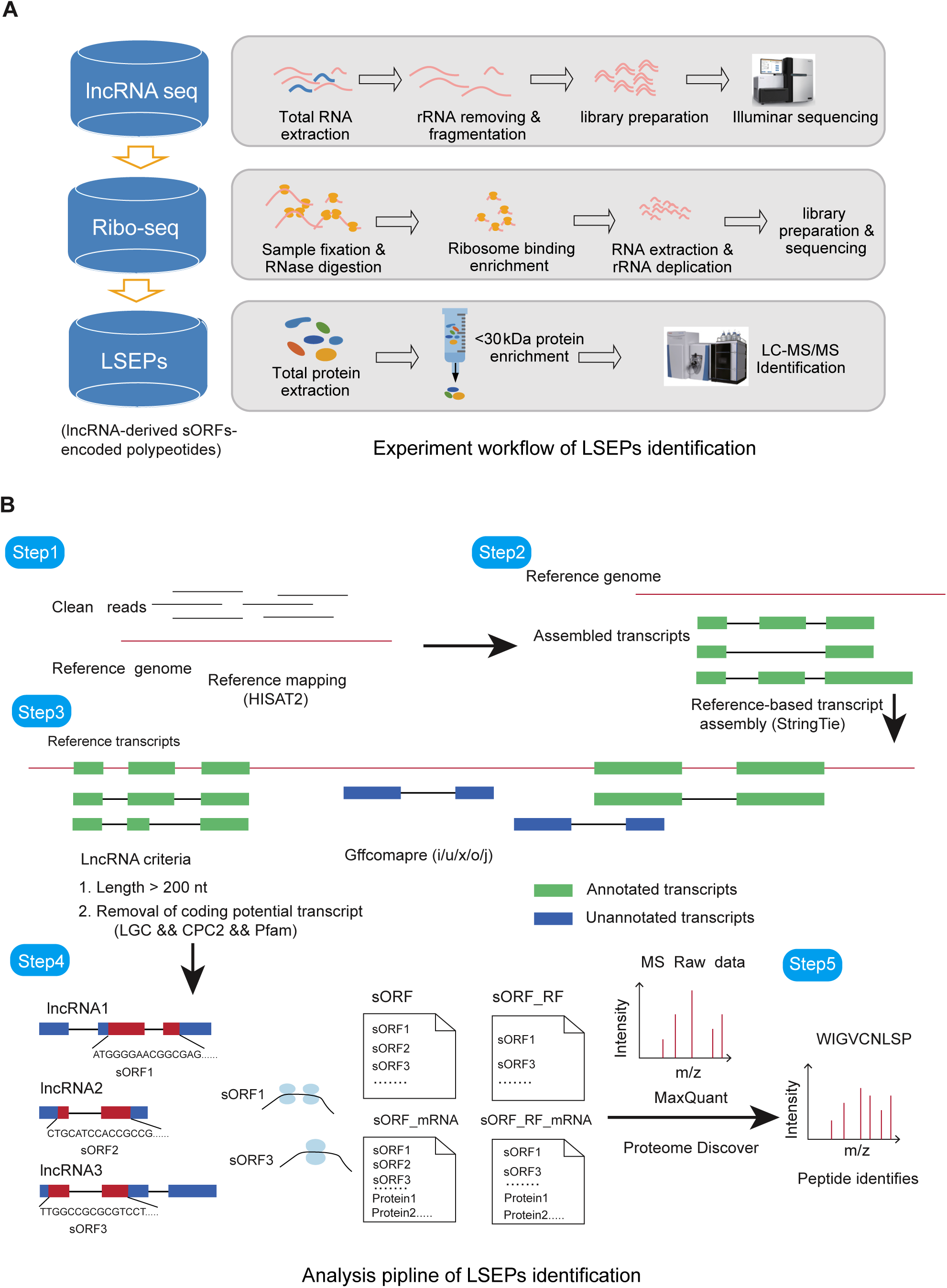
**Workflow and data analysis pipeline of long non-coding RNA-derived sORFs-encoded polypeotides (LSEPs) identification.** (A) Experimental workflow for LSEPs identification. For LncRNA-seq, the total RNA was extracted followed by rRNA removing and RNA fragmentation. The fragmented RNAs was used for the establish of library for Illuminar sequnencing. For Ribo-seq analysis, the fixed samples were digested by RNase, and the ribosome bounded RNAs were enriched and followed by RNA extraction and rRNA removal. After library preparation, the samples were applied for sequencing. For peptide detection, the proteins were extracted and the proteins/peptides < 30 kDa was enriched, and then applied for LC-MS/MS identification. (B) The analysis pipeline for LSEPs identification. The whole genome lncRNAs were identified by standard strand-specific RNA-seq analysis, and those with a length greater than 200 bp were retained. By using CPC2 (0.1), LGC (1.0) and Pfam evaluated the coding potential of lncRNAs. For those lncRNAs with strong coding potential were removed and only lncRNAs that in all three were retained. The getorf tool was used to predict open reading frames in lncRNAs and four small peptide amino acid search databases were constructed based on the presence or absence of ribosomal binding footprints in the open reading frames. These databases included small peptide amino acid search database (sORF), only ribosomal binding small peptide amino acid search database (sORF_RF), small peptide amino acid and rice mRNA amino acid search database (sORF_mRNA), as well as ribosomal binding small peptide amino acid and mRNA amino acid search database (sORF_RF_mRNA). MaxQuant and PD were used to identify translatable small peptides based on four established search databases.

However, it may be possible that ribosome-bound RNAs may not be successful translated. A proteomics-based detection would provide direct evidence of peptide encoding ability. Since limited information on small peptide extraction methods has been reported, we intially tested the extraction buffer preference for rice leaves. The rice leaf samples were homogenized in two different buffers frequently used for plant protein extraction with slight modifications: Buffer A (0.7 M Sucrose, 0.5 M Tris-base, 50 mM EDTA, 10 mM KCl and 2% β-mercaptoethanol) (Lin and Wang, 2014), and Buffer B (10% Glycerol, 50 mM Tris-Cl pH 7.6, 5 mM EDTA, 5 mM EGTA, 2 mM DTT) (Lin et al., 2020). It has been reported that hydro chloride (HCl) benefits the extraction of small peptides in animal (Ma et al., 2016). Therefore, additional HCl with a gradient concentration was used to test the peptide extraction efficiency. After total protein extraction, the small peptides were further enriched by filtering with a 30 kD centrifugation filter and analyzed on SDS-PAGE. As we have shown, both Buffer A and Buffer B are sufficient for small peptide extraction (**Fig. 2**). Furthermore, the additional HCl with a concerntration of 100 mM significantly benefits for the enrichment of small peptides during protein extraction (**Fig. 2**).

**Figure 2.**
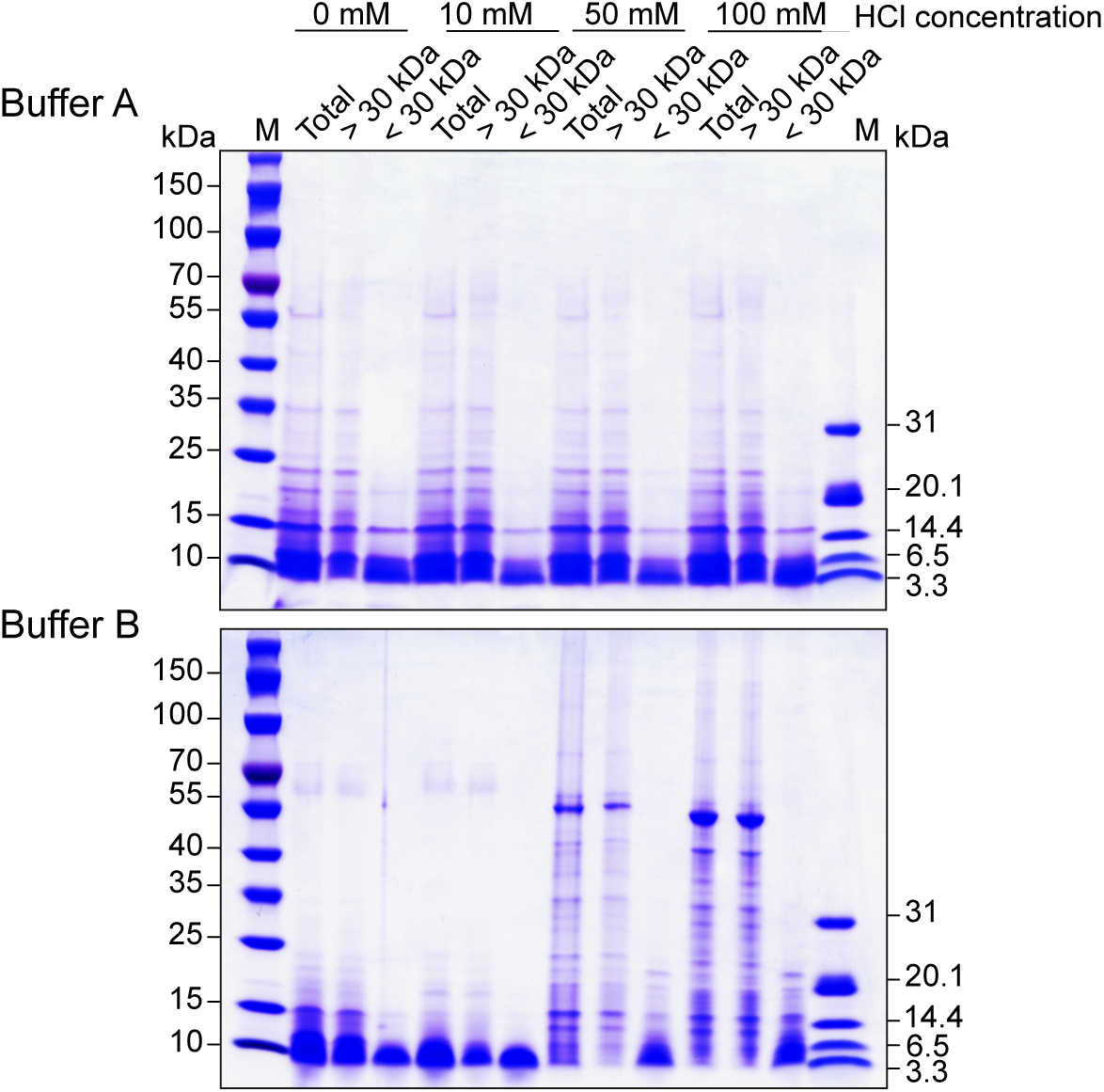
**Protein sample preparation for LC-MS/MS analysis.** Total proteins from rice leaves were extracted using two different extraction buffers (Buffer A and Buffer B) each supplemented with additional HCl at indicated concerntration. The extracted proteins were filtered through 30 kDa ultracentrifuge filters to separate the proteins based on their molecular weight. The upper layer (> 30 kDa) and flow through (< 30 kDa) were then collected, separated on SDS-PAGE, and stained with Coomassie Brilliant Blue for visualization.

The integrity of the searching database significantly affects peptide detection for mass spectrometry identification. Therefore, we established an LSEPs search database based on our lncRNA-seq and Ribo-seq results, as outlined in **Fig. 1B**. Briefly, novel transcripts with length > 200 nt were collected after *de novo* mapping and assembly of lncRNAs. Transcripts with known coding sequences were then eliminated to detect novel peptides derived from lncRNAs. The nucleotide sequence of lncRNAs with small ORFs ranging from 15 to 450 nt were translated into amino acid sequences using the NCBI genetic code table1 (including alternative initiation codons ATG/CTG/TTG), and four custom peptide search databases were constructed. Predicted lncRNA-encoded sORFs with a start codons of AUG, CUG, or UUG within a restricted length range from 15 nt to 450 nt, with or without knowing coding mRNAs, were used to build the sORF (64,730 sequences) and sORF_mRNA (sORF and mRNA encoding sequence, 131,068 sequences) databases, respectively. Similarly, databases for detected ribosome-bound small peptides with or without mRNAs were built as sORF_RF (sORF sequence with ribosome binding, 8,996 sequences) and sORF_RF_mRNA (sORF with ribosome binding and mRNA encoding sequence, 75,334 sequences), respectively. Two frequently used software packages, MaxQuant (Max-Planck-Institute of Biochemistray) and Proteome Discoverer (PD) (Thermo Fisher Scientific), were employed for peptide identification using these four constructed small peptide databases.

### Distribution of identified small peptides in PD and MaxQuant

Based on RNA-seq datasets, a total of 2,764 lncRNAs were identified and categorized into six distinct classes according to their chromosomal positioning: sense lncRNA, intergenic lncRNA, antisense lncRNA, intronic lncRNA, bidirectional lncRNA, and promoter lncRNA (**Fig. 3A**). Our results revealed that the largest proportion (34%) of lncRNAs were located in the intergenic regions, followed by those within gene bodies (21%) and in antisense regions of genes (22%) (**Fig. 3B**), which is consistent with previous reports (Choi et al., 2022).

**Figure 3.**
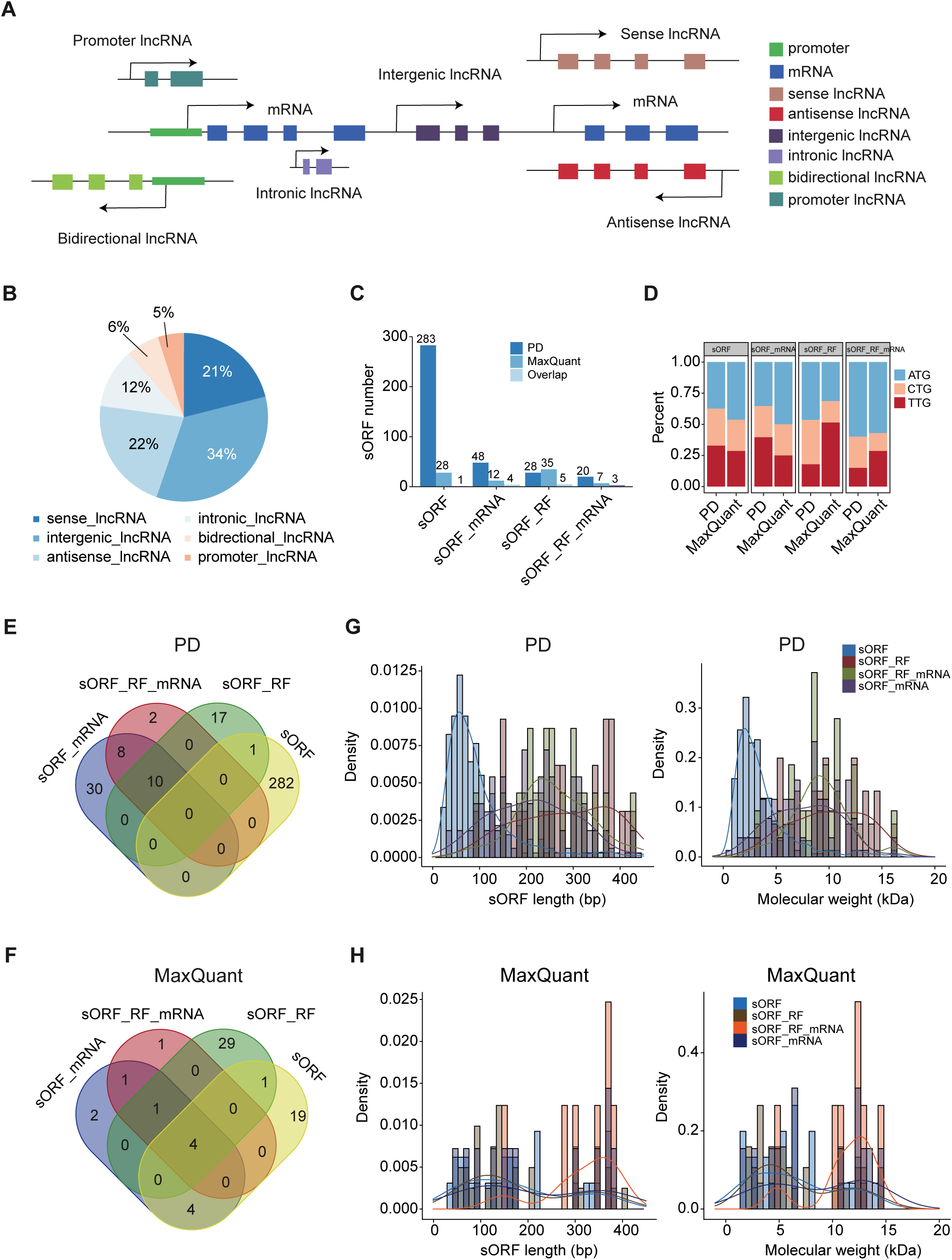
**Overview of translatable open reading frames identified using the LSEPs identification pepline.** (A) Schematic diagram of the location of six types of lncRNAs. (B) The percent of different lncRNA categories identified in rice genomes. (C) The number of small peptides identified in four different search databases using PD and MaxQuant software, with an emphasis on the overlap of peptides identified by both. (D) Percentage of start codons within open reading frames identified across four different search databases using both PD and MaxQuant software. (E) Overlap of translatable small peptides among four different search databases in PD. (F) Overlap of translatable small peptides among four different search databases in MaxQuant. (G-H) Distribution of open reading frames length (left) and amino acid molecular weight (right) of translation small peptides identified in PD (**G**) and MaxQuant (**H**).

We combined the mass spectrometry results of Buffer A and Buffer B with a suitable mass spectrometry search platform and established a database for peptide searches. By utilizing PD and MaxQuant for mass spectrometry data analysis across four constructed peptide databases, we identified 7 to 283 small peptides. PD demonstrated superior performance in identifying peptides in the sORF, sORF_mRNA, and sORF_RF_mRNA databases compared to MaxQuant (**Fig. 3C**). Notably, the sORF database yielded the highest number of identified peptides using PD, showing a six-to-tenfold superiority to others. While searching for sORFs with mRNA information, as showed in the sORFs_mRNA database, this number dramatically dropped, suggesting that the sequences of the coding portion greatly influence the identification of LSEPs. However, this phenomenon not observed between the sORF_RF and sORF_RF_mRNA databases, indicating that our translatome data is quite valuable for LSEPs identification. Conversely, the sORF_RF database showed the greatest peptides yield in MaxQuant, despite having the smallest number of searchable sequences. These results suggest that an optimal database size and a suitable search platform are crucial for peptide detection efficiency. Strikingly, the overlap in peptide identifications between the two tools for any given database was minimal.

The bias in codon usage is a significant evolutionary characteristic of a genome, offering valuable insights into gene functionality and expression patterns. We examined the codon usage bias for the identified LSEPs by predicting the start and stop codons, as previously reported (Zhang et al., 2021). Concisely, an in-frame AUG or a closely matching codon situated within a Kozak context was presumed to denote the start codon. Otherwise, the initiation codon for LSEPs was considered undetermined. Moreover, searches with different databases displayed varying preferences. For example, the sORF_RF_mRNA database favored AUG in both PD (60%) and MaxQuant (57%). Conversely, preferences diverged in PD and MaxQuant for the sORF_RF and sORF_mRNA databases, with AUG and UUG dominance switching between the tools. The sORF database lacked a clear preference in PD but favored AUG in MaxQuant (**Fig. 3D**), supporting prior evidence of prevalent non-AUG initiation in small peptides (Slavoff et al., 2013). Given the importance of Kozak sequence features for translation initiation (Kozak, 1986, 2002), we analyzed motifs surrounding the start codons of identified sORFs and classical mORFs. The classic mORF start codons were accompanied by typical Kozak sequences, whereas those sORFs, regardless of the start codon type, showed marked deviations (**Supplementary** Fig. 2), implying distinct translation recognition mechanisms for small peptides in rice.

An overlap analysis of identified peptides across databases revealed a few peptides consistently detected across multiple databases in both PD and MaxQuant (**Fig. 3E and 3F**). Furthermore, the identified peptides displayed variable distributions in nucleotide length and molecular weight across databases. In PD, sORF-derived peptides predominantly fell below 100 bp and 5 kDa, whereas peptides from other databases were mostly above these thresholds (**Fig. 3G**). Conversely, MaxQuant revealed a contrasting pattern, with sORF_RF_mRNA peptides typically exceeding 200 bp and 10 kDa, unlike the more uniform distribution seen in the remaining databases (**Fig. 3H**). These discrepancies suggest that current analytical tools may have limitations in fully uncovering the small peptides landscape within our databases.

### Correlation between lncRNAs and small peptides

Using mass spectrometry database search results, we identified 62 small peptides through MaxQuant and 350 through PD, across four databases. These peptides mapped to 60 and 316 distinct lncRNAs, respectively. In the PD dataset, lncRNAs were categorized with a notable prevalence of sense lncRNA (31%), followed by intergenic lncRNA (29%) and antisense lncRNA (23%). A similar distribution pattern emerged in the MaxQuant analysis, with a slight increase in sense lncRNA and a minor reduction in antisense lncRNA proportions (**Fig. 4A**). Intriguingly, when assessing lncRNAs expression levels using FPKM values, we found that lncRNAs capable of translating small peptides exhibited significantly higher expression compared to those that did not (**Fig. 4B**). This implies a positive correlation between lncRNA expression and peptides translation potential, a finding consistent with prior animal studies (Aspden et al., 2014; Zeng et al., 2018). Differential analysis of lncRNA types revealed that intergenic and antisense lncRNAs displayed both heightened expression and a greater number of identified small peptides. Conversely, promoter lncRNAs showed high expression despite fewer peptides being identified, and sense lncRNAs, despite being the least expressed, yielded the highest peptide count. This suggests a complex regulatory mechanism in lncRNA-sORFs translation. Notably, only antisense, intergenic, and intronic lncRNAs associated with small peptides displayed consistently elevated expression relative to their respective lncRNA categories, hinting at a possible link to lncRNAs conservation (**Fig. 4C-4F**).

**Figure 4.**
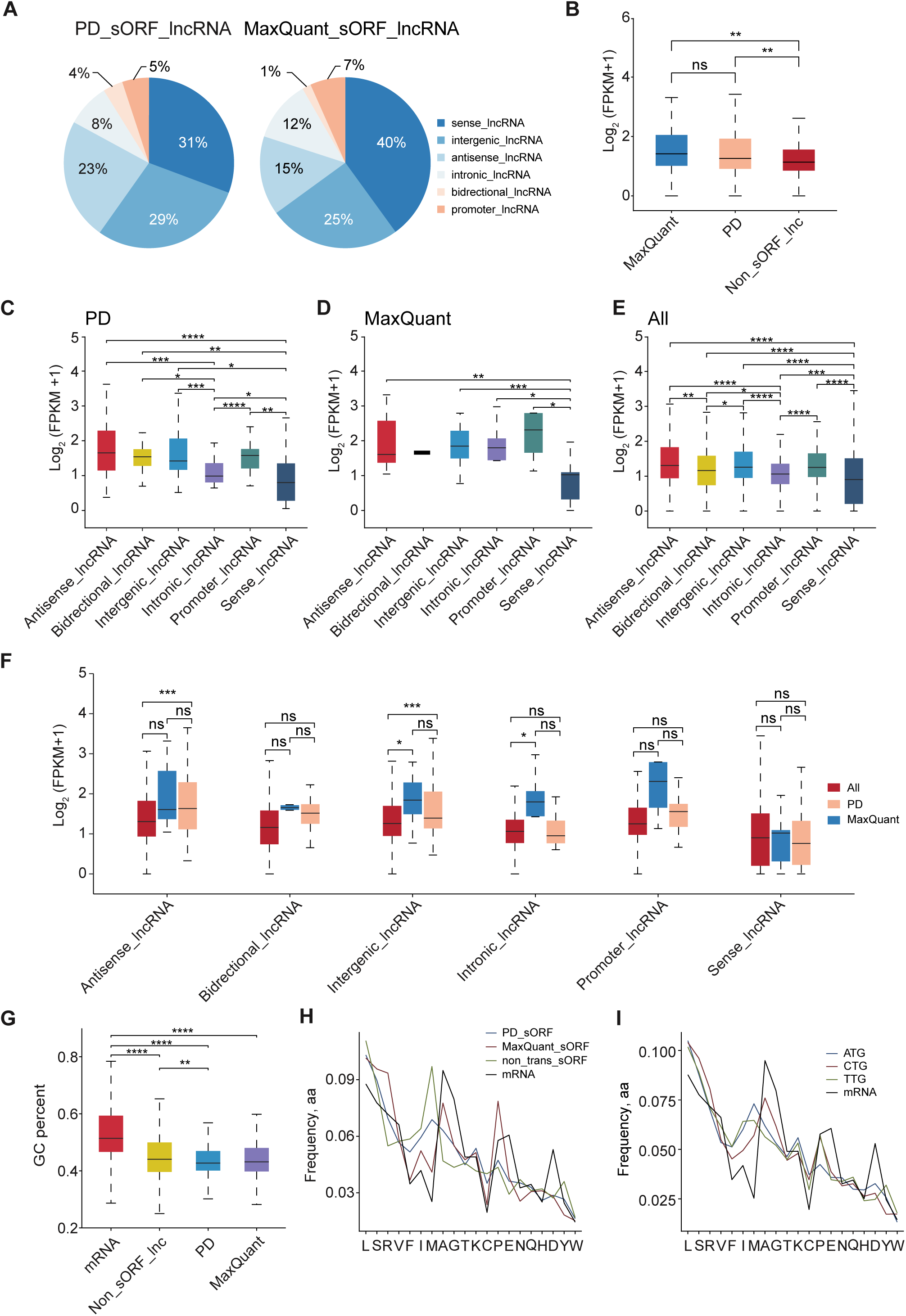
**Characteristics of translatable lncRNAs.** (A) Percentage of six categories of translatable lncRNAs identified in PD (left) and MaxQuant (right). (B) Box plot of lncRNAs expression level, divided lncRNAs into three categories based on the presence or absence of translatable small peptides are the tool used for identification (MaxQuant or PD). (ns and ** respectively indicate p > 0.05 and p < 0.01, Wilcox-test). (C-D) Box plot of expression levels for six categories of translatable lncRNAs identified in PD (C) and MaxQuant (D). (E) Box plot of expression levels for six categories of all lncRNAs. (F) Box plot of expression levels for six categories of lncRNAs. The three colours represent all lncRNAs, translatable lncRNAs identified by PD, and translatable lncRNAs identified by MaxQuant, respectively. (ns, *, and *** respectively indicate p > 0.05, p < 0.05 and p < 0.001, Wilcox-test). (G) Boxplot plot showing the GC content percentage of mRNAs, lncRNAs without identified small peptides by PD and MaxQuant (Non_sORF_lnc), translatable lncRNAs identified in PD (PD), and translatable lncRNAs identified in MaxQuant (MaxQuant). (H) Comparison of amino acid compositions among mRNAs, identified small peptides in PD and MaxQuant, and other predicted small peptides that were not been identified (Non_trans_sORF). (I) Comparison of amino acid compositions of mRNAs and identified small peptides with three different stat codons.

In our exploration of transcript features predictive of translation, we observed that lncRNAs encoding small peptides had a significantly lower GC content compared to rice mRNAs (**Fig. 4G**). This finding echoes previous studies that have implicated GC content and sequence conservation in translation potential (Ulitsky, 2016; Choi et al., 2019; Ruiz-Orera and Albà, 2019). Our dataset further revealed differences in amino acid composition between the small peptides and functionally annotated proteins. Notably, the frequencies of the hydrophobic amino acids methionine (M) and isoleucine (I) frequencies were higher, while the frequencies of alanine (A), glutamic acid (E), and aspartic acid (D) were lower (**Fig. 4H**). This resonates with sORF profiles reported in mammalian lncRNAs (Couso and Patraquim, 2017). Strikingly, this compositional disparity persisted even when comparing unidentified versus identified small peptides. This suggests that the limitations lie in detection methodologies rather than true biological variation (**Fig. 4H**). While investigating the influence of start codons, we found no discernible differences in amino acid profiles among small peptides initiated by AUG, CUG, or UUG. This reinforces the uniformity in translation initiation across these codons (**Fig. 4I**).

### Distribution of small peptides in the rice genome

To investigate the genomic distribution of small peptides, we employed a 100-kilobase pair (kb) sliding window strategy across each rice chromosome, counting the number of peptide number per window. Our findings unveiled a strikingly non-uniform distribution of small peptides across the chromosomal expanse (**Fig. 5A**). For each peptide, we calculated the proximity to centromeres and found a conspicuous enrichment of 31 (8.86%) small peptides in the centromeres region in the PD search. Conversely, the MaxQuant search showed a more uniform chromosomal dispersion, which may be attributed to the sparse peptide detection and weaker statistical representation (**Fig. 5B**).

**Figure 5.**
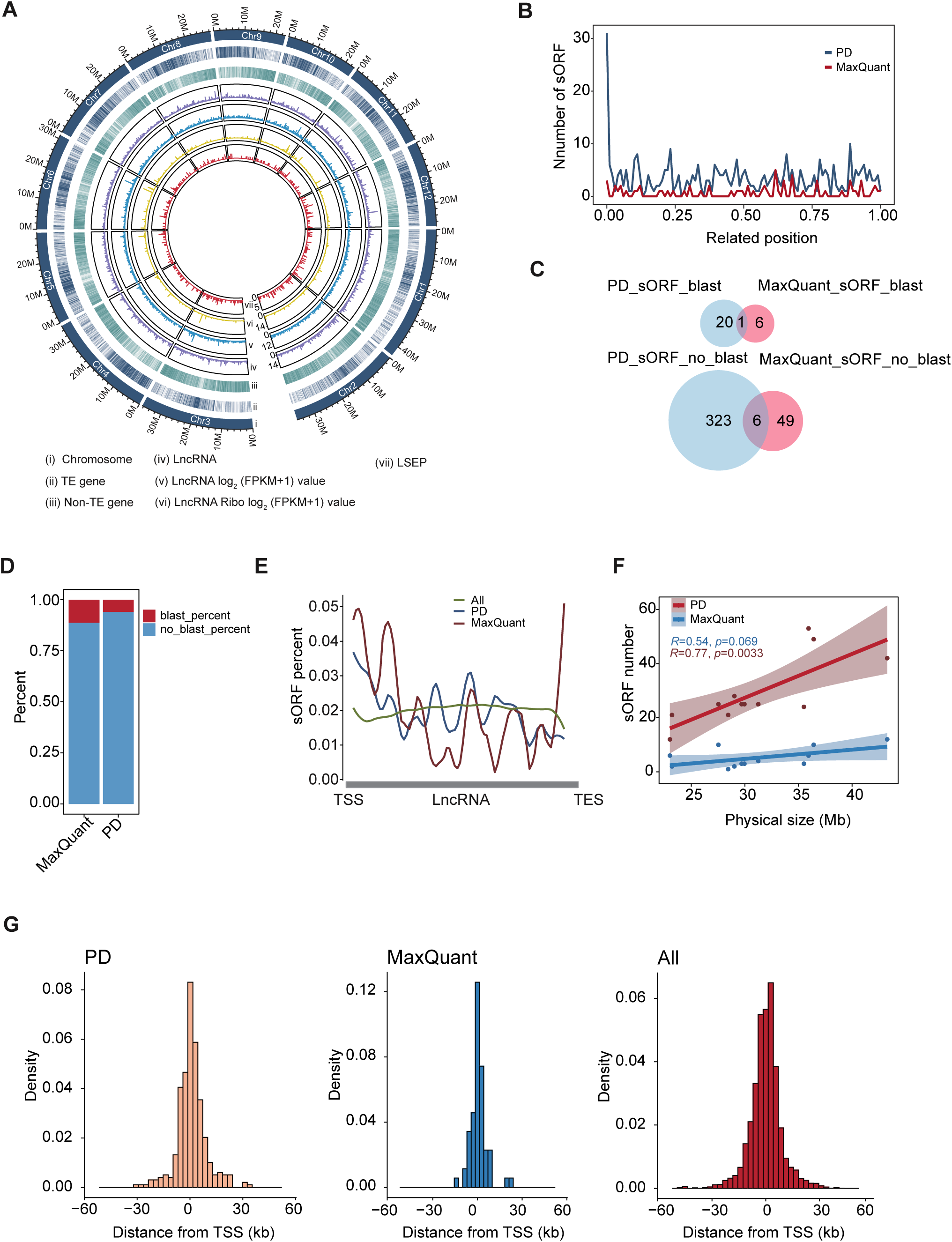
**Distribution characteristics of small peptides on the rice genome.** (A) Genome-wide distribution of transposable element (TE)-gene, Non-TE gene, small peptides (LSEPs), LncRNA, and their expression level. (B) Distributions of small peptides identified by MaxQuant (red) and PD (blue) along the chromosomal arms in 10 kb sliding window. The x-axis represents the normalized length of each arm with centrome set to ‘0’ and the telomere to ‘1’. (C) The number of overlapping small peptides identified by MaxQuant and PD that can be aligned to rice mRNA (top). The number of overlapping small peptides identified by MaxQuant and PD that cannot be aligned to rice mRNA (bottom). (D) Bar chart showing the percentage of translatable small peptides identified in MaxQuant and PD that can be aligned to rice mRNA (red) or cannot be aligned to rice mRNA (blue). (E) Distribution of translatable (blue, red) and all (green) small open reading frames on lncRNAs. All lncRNAs were scaled to the same length. Blue: lncRNAs of translatable small peptides identified by PD; red: lncRNAs of translatable small peptides identified by MaxQuant; green: all identified lncRNAs; TSS, transcription start site; TES, transcription end site. (F) Scatter plot showing the correlation between chromosome physical length and the number of small peptides identified by MaxQuant (blue) and PD (red). (G) Histograms showing the distance from each small open reading frames to the TSS of the closest mRNA. Left: small peptides identified by PD; middle: small peptides identified by MaxQuant; right: all predicted small open reading frames.

Given the exclusion of rice mRNA sequences in the sORF and sORF_RF databases that might lead to misidentification, we used BLASTP to map peptides from these databases onto rice mRNA amino acids(Camacho et al., 2009). Strikingly, only a small fraction of peptides aligned successfully: 21 (6%) in MaxQuant and 7 (11.3%) in PD, with negligible overlap between datasets – a single peptide was identified in both platforms, and the remaining peptides showed minimal coincidence (**Fig. 5C and 5D**). These observations underscore the low likelihood of identified peptides being mere fragments of mRNA amino acids and emphasize the unique profiling capabilities of MaxQuant and PD.

To further expand our understanding of the distribution of sORFs within lncRNAs, we normalized lncRNAs length and discerned a remarkable enrichment of small peptides near transcription start sites (TSSs) in both PD and MaxQuant searches, with an additional enrichment at transcription end sites (TESs) detected uniquely in PD (**Fig. 5E**). Furthermore, consisted with previous reports, there is a positive correlation between the number of identified LSEPs per chromosome and the chromosome length (Wang et al., 2020; Pei et al., 2022) (**Fig. 5F**).

It is well known that lncRNAs usually play roles in regulating neighboring genes. We then calculated the distance of each LSEP to the nearest mRNA TSS to explore the spatial relationship between LSEPs and neighboring genes. Notebly, 15.7% and 18.3% of LSEPs in PD and all predicted, respectively, resided beyond 10 kb from the closets gene’s TSS, starkly contrasting with only 4.8% in MaxQuant (**Fig. 5G**). Collectively, these findings illustrate the broad dispersion of LSEPs in the genome, avoiding active transcription regions, and highlight the differential efficiency of MaxQuant and PD in LSEPs identification.

### Characteristics of small peptides in coding and non-coding region

In our investigation of all recognized LSEPs using MaxQuant and PD, we noticed that more than half of the detected small peptides were located in the forward strand, with proportions standing of 181 (53.2%) via MaxQuant and 33 (51.7%) via PD (**Fig. 6A**). After categorizing these peptides based on their genomic positions, a substantial proportion, 56.3% in PD and 43.5% in MaxQuant, were found within intergenic regions (**Fig. 6B** and **Fig. 6C**), suggesting a high translational potential for intergenic sequences. Furthermore, by assessing the number of LSEPs in each lncRNA, we evaluated their translational potential and observed that multiple translated small peptides within a single transcript predominantly occurred in antisense and sense lncRNAs (**Fig. 6D** and **Fig. 6E**). This indicates that these types of lncRNAs exhibit stronger translational capabilities than other translatable lncRNA types.

**Figure 6.**
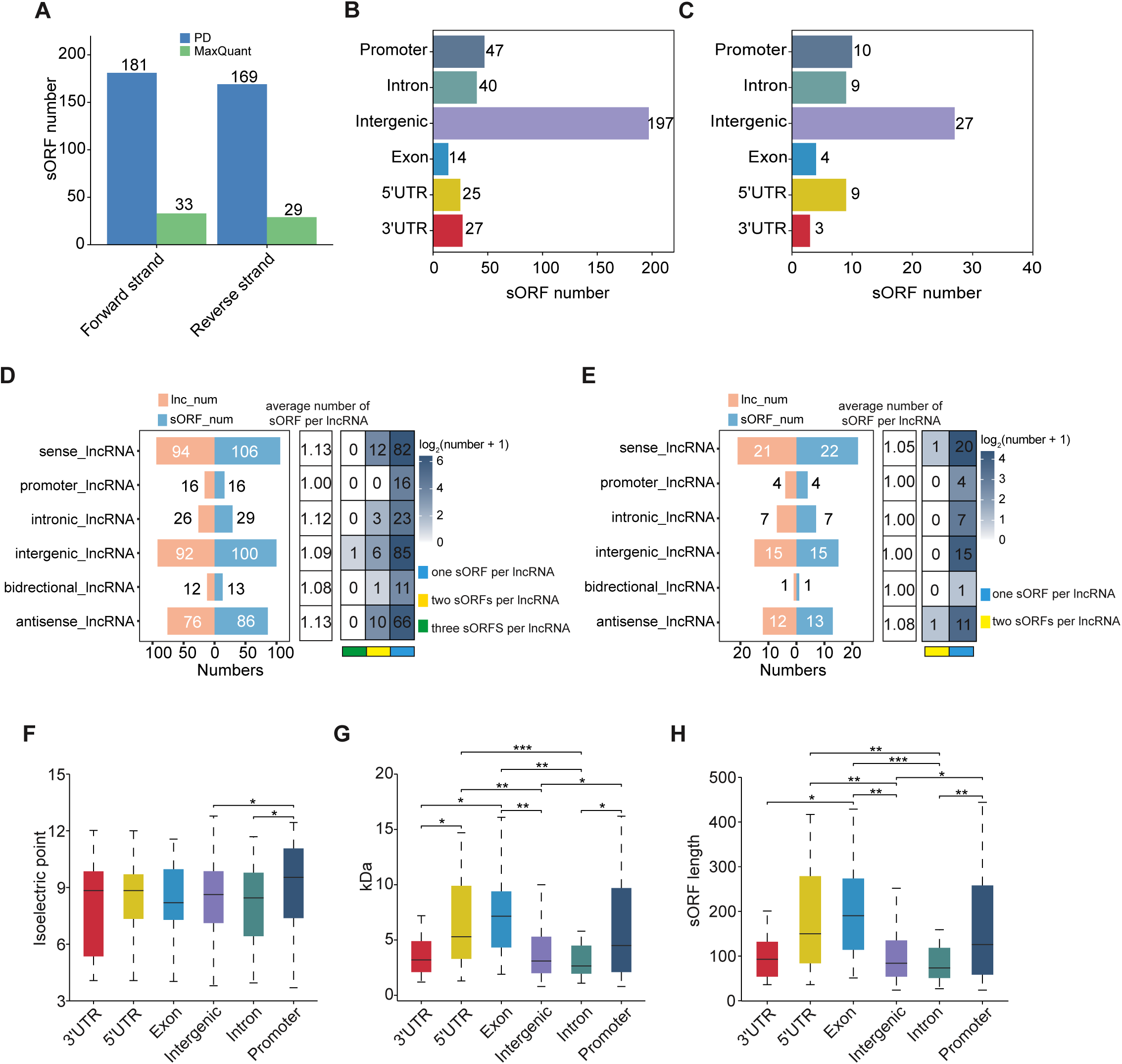
**Characteristics of identified small peptides.** (A) The number of small peptides identified by MaxQuant and PD derived from forward and reverse strands. (B-C) The number of small peptides identified by PD (B) and MaxQuant (C) in different region of the rice genome. (D-E) Statistics of the number of small peptides in all types of lncRNAs for PD (D) and MaxQuant (E). (F-G) The isoelectric point (F) and molecular weight (G) of small peptides in different regions of the rice genome. The Wilcox-test was used for hypothesis testing. ns: p > 0.05; *: p < 0.05, **: p < 0.01, ***: p < 0.001. (H) The length of translatable small open reading frames derived from different regions of the rice genome. The Wilcox-test was used for hypothesis testing. ns: p > 0.05; *: p < 0.05, **: p < 0.01, ***: p < 0.001.

To assess the efficiency between Buffer A and Buffer B, we collected quantitative data from all proteomic databases and found that Buffer B generally outperformed in capturing small peptide abundances, specifically in PD searches. Buffer B yielded higher abundance for small peptides across databases. For PD, 10.4% and 64.6% of small peptide abundances were greater than 10 in buffer A and buffer B for the sORF_mRNA database, 20% and 85% for sORF_RF_mRNA database, 21.4% and 82.1% for sORF_RF database, 55.5% and 59.4% for the sORF database (**Supplementary** Fig. 3A). We also found that the abundance of small peptides in the 3’UTR region identified by buffer B was higher than that of buffer A (**Supplementary** Fig. 3C). Parallel assessments on mRNA and results of MaxQuant confirmed a similar pattern, with Buffer B demonstrating superior performance in capturing mRNAs and LSEPs abundances across sORF-related databases (**Supplementary** Fig. 3B, 4A, and **4B**). This affirms Buffer B’s overall superiority for both small peptide and mRNA analyses.

In probing the distinct properties of LSEPs, we conducted a comprehensive analysis by calculating key physicochemical attributes, such as isoelectric points, molecular weights, and lengths. Notably, peptides originating from promoter regions exhibit the highest isoelectric points. Moreover, peptides in the 3’UTR, intergenic, and intronic regions display significantly reduced molecular weights and lengths relative to those in other genomic compartments, a pattern consistent across MaxQuant and PD datasets (**Fig. 6F-6H** and **Supplementary** Fig. 4C and 4D). Collectively, these findings accentuate the distinct characteristics of LSEPs derived from different genomic landscapes, highlighting the importance of considering regional context when studying these bioactive molecules.

### Association between epigenetic markers and translatable lncRNAs

Epigenetic mechanism involved in plant stress response and development are well characterized, primarily through the regulation of protein-coding genes expression (Chang et al., 2020; Lloyd and Lister, 2022). Given our observation that small peptides show a preference for appearing on highly expressed lncRNAs, and considering previous reports that epigenetic markers regulate lncRNA expression (Hung et al., 2020; Zhao et al., 2021), we explored the relationship between translatable lncRNAs and epigenetic markers. We investigated the distribution of epigenetic markers within 2 kb upstream and downstream of lncRNAs, including DNase I hypersensitive sites (DH), histone H3 lysine 36 trimethylation (H3K36me3), H3K4me3, H3K9 acetylation (H3K9ac), H3K27me3, H3K9me2. We found that the signal of activation markers at the TSS site of translatable lncRNAs was higher than that of overall and untranslated lncRNAs. In contrast, the signal of the represive marker H3K9me2 in the gene body region of translatable lncRNAs was lower than that of the overall and untranslated lncRNAs (**Fig. 7A**).

**Figure 7.**
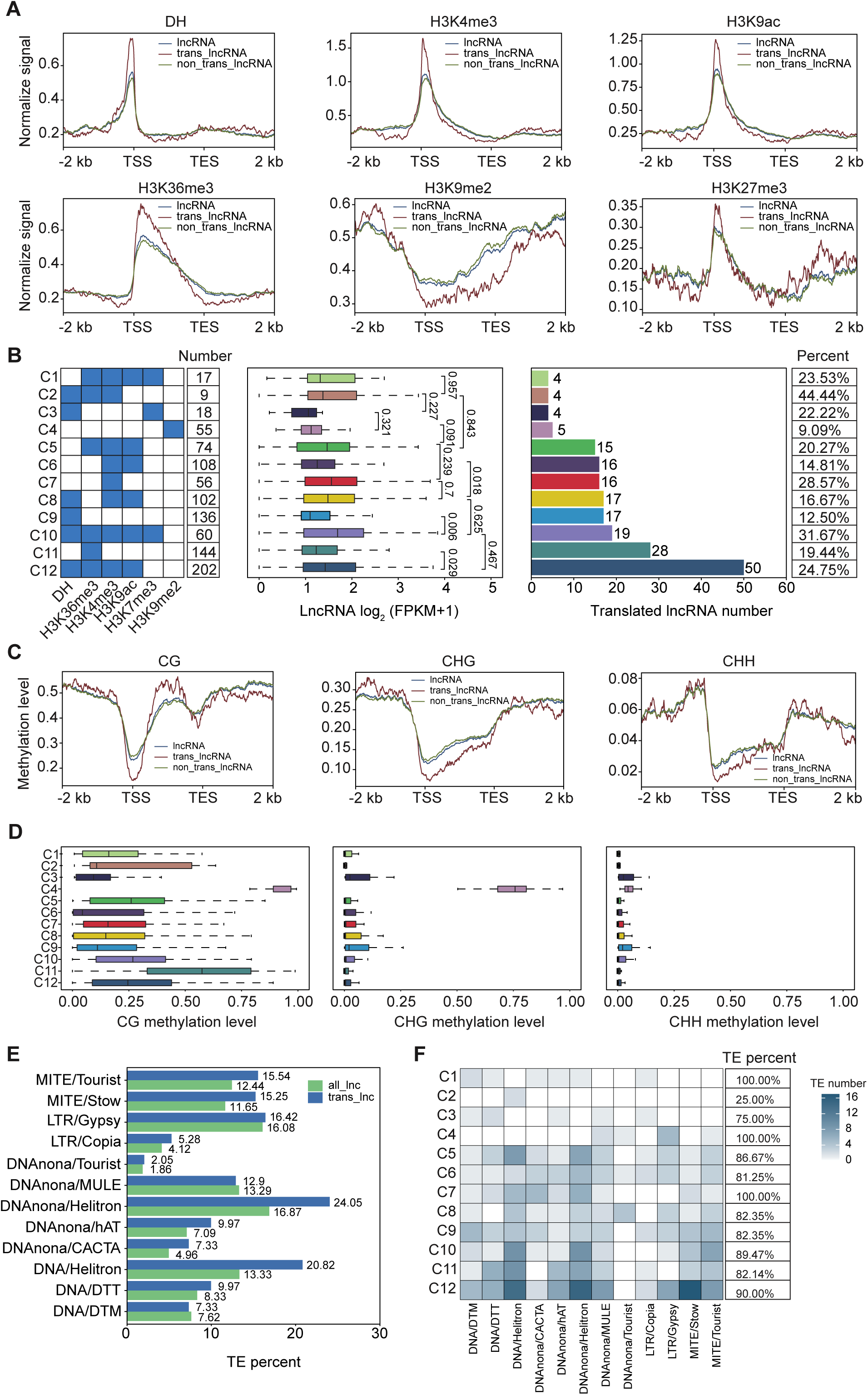
**The association between translatable lncRNAs and DH, epigenetic markers and TE.** (A) Distribution of DH and histone modification signal intensity in the overall lncRNAs, translatable lncRNAs, untranslated lncRNAs, and their 2 kb up/down flanking regions. TSS, transcription start site; TES, transcription end site. (B) The number of 12 types of epigenetic related lncRNAs, box plot of their expression level, bar chart of the number of translatable lncRNAs, and percentage of translatable lncRNAs. (C) Distribution of CG, CHG and CHH methylatioin level in the overall lncRNAs, translatable lncRNAs, untranslated lncRNAs, and their 2 kb up/down flanking regions. TSS, transcription start site; TES, transcription end site. (D) Box plot of CG, CHG and CHH methylation level of 12 type epigenetic related lncRNAs. (E) Bar chart of the percentage of TE-related lncRNAs in overall lncRNAs and translated lncRNAs. (F) The number and percentage of 12 types of epigenetic related lncRNAs in the TE regions.

We conducted a statistical analysis of lncRNAs with epigenetic marks and found that the proportion of translatable lncRNA regions containing activating marks is higher compared to overall lncRNAs. Conversely, the proportion of the repressive mark H3K9me2 is lower in translatable lncRNAs. Specifically, translatable lncRNA regions have the highest proportion of the activating mark H3K4me3, while the proportion of the repressive mark H3K9me2 is the lowest (**Supplementary** Fig. 5A). To further determine if epigenetic markers can regulate lncRNA expression similarly to mRNA, we investigated the relationship between these markers and lncRNAs expression level. We found that lncRNAs with activation markers had significantly higher expression level than those without such markers. For repressive markers, there was no significant change, which is likely due to other regulatory mechanisms of lncRNAs (**Supplementary** Fig. 5B). Additionally, we divided lncRNAs (FPKM ≧ 1) into three categories based on their expression level from high to low. We discovered that highly expressed lncRNAs often had higher activation markers near their transcription start sites, while gene body regions often had lower repressive markers (**Supplementary** Fig. 5C). These results suggest that translatable lncRNAs are more likely to be located in regions with epigenetic activation markers, and their expression is regulated by these markers. This feature may also suggest that translated peptides have complex regulatory mechanisms involved in plant growth and development.

Considering that multiple epigenetic markers often co-regulate the expression of protein-coding genes, we classified lncRNAs into 12 types based on the locations of multiple epigenetic markers. We found that lncRNAs containing multiple activation marker (C5, C6, C8, C10, C12) tended to have more translatable lncRNAs and higher expression level. However, only a few of these lncRNAs showed significant differences in expression levels. For lncRNAs (C4) containing only H3K9me2 repressive markers, the number and proportion of translatable lncRNAs were lower. These results indicate that the expression of lncRNAs is regulated by epigenetic markers, although the degree of synergistic regulation by multiple markers is limited. Moreover, translatable lncRNAs predominantly occur in regions with activation markers (**Fig. 7B**).

DNA methylation is another crucial epigenetic marker that regulates the expression of protein-coding genes (Zhang et al., 2018b; He et al., 2022). We compared the distribution of DNA methylation across lncRNAs and found that CG methylation was most prevalent in the gene body region of translatable lncRNAs, followed by overall lncRNAs and untranslated lncRNAs. In contrast, the distribution of CHG and CHH methylation was exactly opposite of that observed for CG methylation (**Fig. 7C**). Similarly, we categorized lncRNAs (FPKM ≧ 1) into three groups based on their expression level, from high to low, to explore the relationship between DNA methylation and lncRNAs expression. For CG methylation, we found that the methylation level in the gene body region did not follow the expected pattern of distribution according to expression levels. In contrast, CHG and CHH methylation levels were generally lower in high-expression lncRNAs and higher in low-expression lncRNAs (**Supplementary** Fig. 5D). These results suggested that DNA methylation can regulate lncRNA expression to some extent, with translated lncRNA regions typically showing higher CG methylation and lower CHG and CHH methylation. To further understand the coordinated regulation of DNA methylation and histone modifications, we measured DNA methylation level across 12 types of lncRNA regions. As expected, lncRNA regions marked by activating histone modifications (C12, C11, C10) generally exhibit higher CG methylation level and lower CHG and CHH level. Interestingly, these lncRNAs also contain the highest number of translatable lncRNAs. In contrast, for repressive marks, it was surprising to find that C4 (H3K9me2) lncRNA regions exhibit high level of all three types of methylation, while C3 (H3K27me3, DH) lncRNA regions showed results consistent with expectations, with generally lower CG methylation level and higher CHG and CHH methylation level (**Fig. 7D**).

Transposable elements (TEs) are important regulatory elements in plant genomes and are widely distributed in the genomes of both plants and animals. Recent reports have indicated that TE-lncRNAs play a critical role in plant responses to abiotic stress (Wang et al., 2017). Given the significant role of TE-lncRNAs, we compared the number of TEs present in translatable lncRNAs versus overall lncRNAs. We found that the proportion of translatable lncRNAs containing TEs (81.82%) is higher than that of overall lncRNAs (74.09%) (**Fig. 7E**). Among these, DNAnona/Helitron and DNA/Helitron are the two most prevalent TE types in translated lncRNAs, and they are also more common than in overall lncRNAs. In contrast, DNAnona/MULE and DNA/DTM are less prevalent in translated lncRNAs compared to overall lncRNAs (**Fig. 7E**). To further explore the association between TE-related lncRNAs and epigenetic markers, we counted the number of TE-related lncRNAs across 12 types of lncRNA intervals. We found that TE-related lncRNAs constitute more than 75% in most categories of translatable lncRNAs, with only C2 at 25%. Additionally, DNAnona/Helitron, DNA/Helitron, and MITE/Stow were the most prevalent TE types (**Fig. 7F**). These results suggested a close association between TE-related lncRNAs and translatable lncRNAs, particularly with DNAnona/Helitron and DNA/Helitron TEs. Moreover, the presence of TEs in most epigenetically marked translatable lncRNAs further indicates that translatable lncRNAs may be subject to multifaceted regulation.

### Validation of peptide-encoding potehntial of OsLSEPs

To further validate the accuracy of our multi-omics findings, an experimental strategy was employed to verify the peptide-coding potential of OsLSEPs identified through multi-omics analysis. A random selection of twelve lncRNAs, predicted to encode OsLSEPs with start condons AUG, CUG, and UUG, was made (**Supplementary Table 2A**). A mutated GFP (mGFP), lacking a start codon, was genetically fused to the C-terminal of these lncRNAs and transiently expressed in the epidermial cells of *Nicotiana benthamiana*. Protein expression for nine distinct OsLSEPs was detected using immunoblotting with an anti-GFP antibody and confirmed via confocal microscopy (**Fig. 8A and 8B**). The presence of multiple bands in the immunoblotting assay suggested that certain lncRNAs may encode multiple peptides (**Fig. 8C**). The GFP signals for OsLSEPs were observed in both the nucleus and cytoplasm (**Fig. 8B**), aligning with previous observations in animal cells (Zhang et al., 2021). Three lncRNAs, predicted to have start codon of ATG, CTG, and TTG, respectively, were selected for further validation of their coding regions. The coding region of these lncRNAs were delineated to elucidate the corresponding peptide sequences by excluding the start codons. As demonstrated, the fluorescent signal and peptide accumulation for OsLSEP1m1 and OsLSEP5m1, where the start codons were removed, were eliminated (**Fig. 9A-9D; Supplementary Table 2B**). However, for OsLSEP12, protein accumulation and GFP signal remained unaffected upon removal of the CUG from the sequence (OsLSEP12m1) (**Fig. 9E and 9F; Supplementary Table 2B**). Re-analysis of the OsLSEP12 sequence revealed four potential start codons (**Fig. 9E**). By individually deleting these putative start codons and assessing their expression, we found that only OsLSEP12m3, not OsLSEP12m2 or OsLSEP12m4, abolished peptide expression and GFP signal (**Fig. 9F** and **Supplementary Table 2B**), indicating that the third AUG is essential for OsLSEP12 expression. The constructs were also transiently expressed in rice protoplasts, where OsLSEP1, OsLSEP5, and OsLSEP12 demonstrated peptide-coding capability **(Fig. 9G)**. The removal of start codon also abolished protein expression, consistent with the results in *N. benthamiana*. Collectively, our data illustrate that rice lncRNAs possess the capability to encode peptides. This study has developed an experimental protocol for the detection of LSEPs in rice.

**Figure 8.**
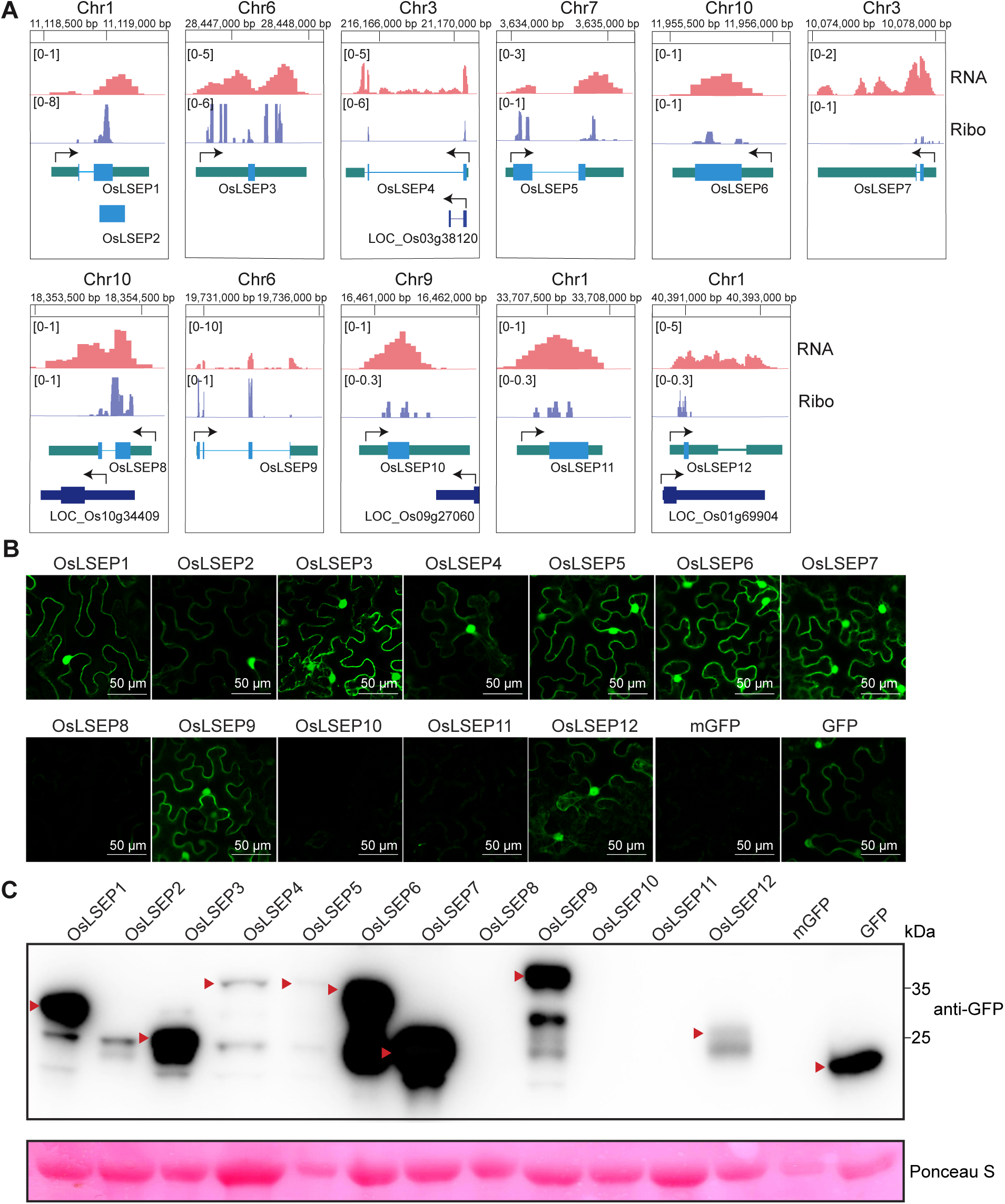
**Peptide encoding validation of identified OsLSEPs.** (A) Diagram of putative OsLSEPs. The normalized reads coverage of 12 selected sORFs in Ribo-seq and RNA-seq is shown. Among them, sORF1, 3, 4 start with ATG; sORF5, 6, 7, 8, 9 start with TTG; sORF2, 10, 11, 12 start with CTG. Gene and lncRNA structure is plotted at the Bottom: light blue box represent the identified small open reading frame, light green box represent lncRNA, dark blue box represent gene. Black arrows indicate direction of transcription of lncRNA and gene. (B) Expression of OsLSEPs in *Nicotiana benthamiana* epidermial cells. The putative OsLSEPs were fused with mutated GFP (mGFP, GFP without start codon) at the C-terminus. GFP flourescence was measured at 24 hours after Agroinfiltration with confocal microscope. mGFP and GFP were used as negative and positive control, respectively. (C) Protein expression of OsLSEPs in *N. benthamiana*. Proteins were extracted from *N. benthamiana* leaves at 24 hours after agroinfiltration. The expressed OsLSEPs were detected by immunoblotting with anti-GFP antibody. Expressed peptides with molecular weight similar with *in silico* predictions are indicated by red arrows. Ponceau S was used as a loading control.

**Figure 9.**
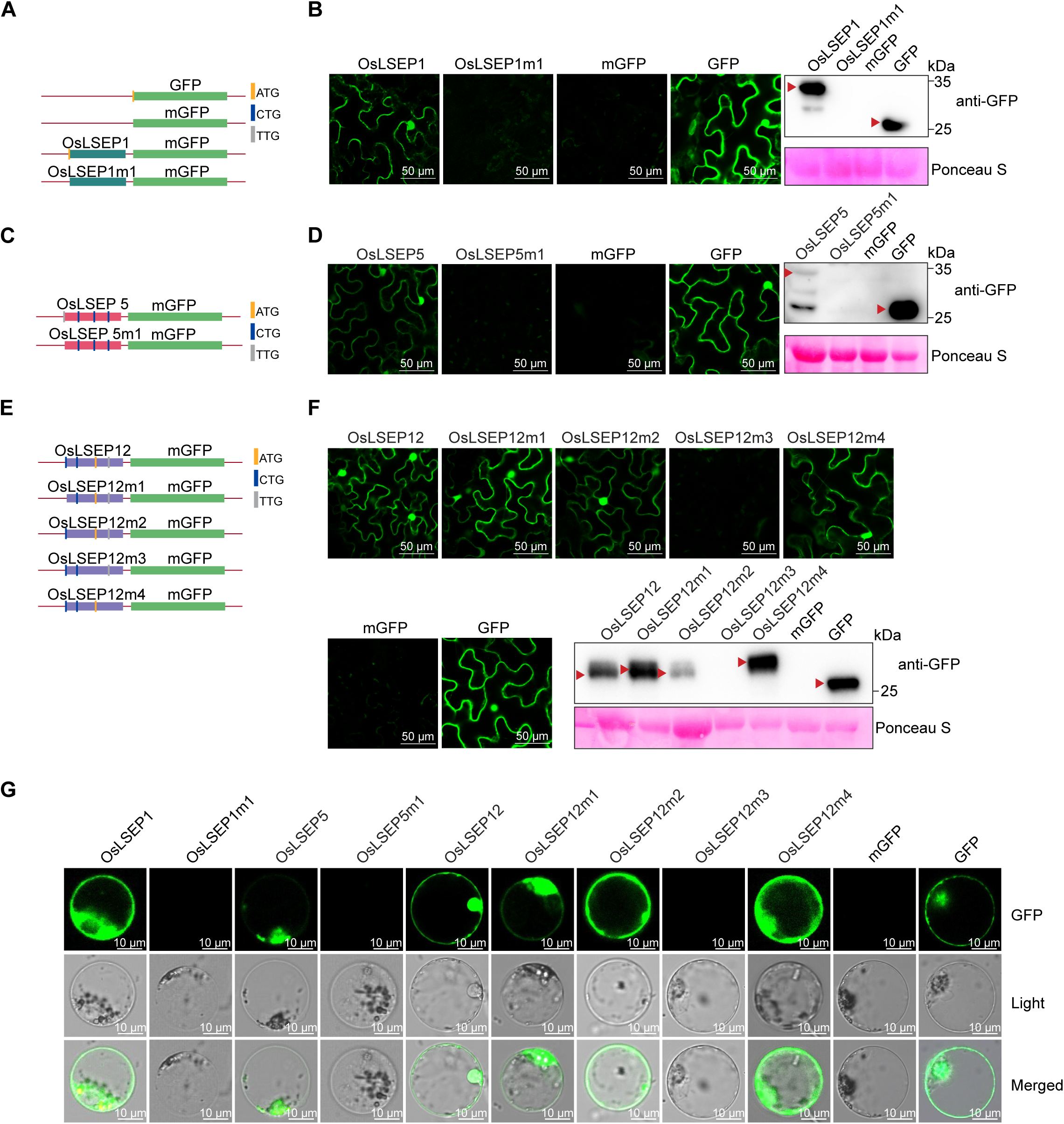
**Peptide identification of OsLSEPs.** (A, C, E) Diagrame of lncRNA ORF cloning strategy. The predicted OsLSEP ORFs and the starting codon-removed ORFs (OsLSEPm) were fused with mutated GFP (mGFP) and transiently expressed in *N. benthamiana* epidermial cells. (B, D, F) Fluorescence of OsLSEP1-mGFP (B), OsLSEP5-mGFP (D), and OsLSEP12-mGFP (F) were detected by confocal microscope. mGFP and GFP were used as negative and positive control, respectively. (G) Fluorescence of OsLSEPs in rice protoplasts. Flouresence signals were measured at 20 hours post-protoplast transformation.

## Discussion

lncRNAs are pervasive in eukaryotic genomes and play pivotal roles in various biological processes. In plants, including Arabidopsis and rice, lncRNAs have been associated with the regulation of flowering time, reproduction, and stress response. They exert their influence by affecting chromatin modification, which in turn impedes the access of regulatory proteins to their intended RNA and DNA targets (Csorba et al., 2014; Shin et al., 2022; Zhang et al., 2023). Interestingly, recent studies have shown that lncRNAs in animals can encode peptides, which are vital for the regulation of numerous biological processes, independent of the intrinsic functions of the lncRNAs themselves (Huang et al., 2017; Tajbakhsh, 2017). However, there is little evidence supporting similar regulatory mechanisms for plant lncRNAs, especially in rice. In this study, we have established an experimental approach to identify polypeptides encoded by sORFs within rice lncRNAs. This method aims to bridge the knowledge gap regarding the potential peptide-coding capabilities of plant lncRNAs and their possible roles in plant biology, paralling the findings in animal systems.

lncRNAs in plants have been implicated in a variety biological processes, including the regulation of flowering time, root development, and immune response (Gao et al., 2020; Huang et al., 2023). These lncRNAs serve as molecular scaffolds, recruiting multiple proteins to specific genomic loci, thereby influencing gene expression. Interestingly, some lncRNAs have been found to encode small peptides, which can further modulate cellular functions. For instance, certain lncRNA-derived peptides may contribute to plant defense mechanisms against pathogens, potentially by affecting the expression of immune-related proteins (Seo et al., 2017; Zhang et al., 2022). In soybean, the *ENOD40* lncRNA encodes two peptides of 12 amino acids and 24 amino acids, which can interact with sucrose synthase to regulate sucrose utilization in nitrogen-fixing nodules of soybeans (Rohrig et al., 2002). In cotton, *GblncRNA7* can encode a small peptide of 75 amino acids, which is similar to PSK3 in Arabidopsis (Zhang et al., 2022). PSK is an important signaling molecule composed of a disulfide-bridged pentapeptide, which can act as a DAMP and was primarily sensed by PSKR1, leading to an increase in cytoplasmic [Ca^2+^] and the activation of auxin-mediated pathways to enhance the immunity (Zhang et al., 2018a; Zhang et al., 2022). Given the significant role of lncRNA-encoded small peptides in plant development and stress responses, there is an urgent need to develop a rapid and high-throughput method for identifying lncRNA-encoded small peptides across the entire plant genome. Recently, by utilizing RNA sequencing and ribosome profiling techniques, 668 lncRNAs with potential peptide coding ability have been identified in rice species (Zhu et al., 2023). However, due to the high potential for false positives in binding assays, it remains unclear whether these lncRNAs indeed encode small peptides. Therefore, protein-level evidence is necessary for the accurate detection of LSEPs. In Arabidopsis, MS-based analysis has led to the identification of small novel peptides (Ohyama et al., 2008). Here, we integrate proteomics approaches with RNA sequencing and ribosome profiling methods, leading to the identification of a total of 403 LSEPs in rice. Furthermore, the peptide coding ability was validated by transient expression in plant, providing experimental evidence that these LSEPs are not partial fragments of longer protein sequences. Together with previous findings, our results provide valuable resources for understanding the biological functions of LSEPs in plants.

We sought to explore the potential mechanisms underlying the regulation of small peptides. Epigenetic marks can respond to environmental change and regulate the expression of mRNAs, playing a crucial role in plant growth and development. In our study, we found that epigenetic marks also have a regulatory effect on the expression of lncRNAs, which is similar to previous reported (Hung et al., 2020). We investigated the association between translatable lncRNAs and epigenetic marks, finding that a greater number of small peptides are identified in regions with activating marks, which may be lncRNAs in these regions have higher expression level, leading to a more abundance of small peptides, making them easier to identify. As mentioned in Figure4B, the expression level of translatable lncRNAs are significantly higher than those of non-translatable lncRNAs. Therefore, further confirmation is needed to determine whether the disappearance and enhancement of activating marks in response to environmental and stress changes will affect the abundance of small peptides within lncRNAs. In Arabidopsis, studies have shown that the HDA6-LDL1/2 histone modification complex can regulate the levels of H3Ac/H3K4me2 in the promoter regions of lncRNAs to mediate lncRNA expression (Hung et al., 2020). In mammals, several lncRNAs regulated by H3K27ac and DNA methylation have been identified to participate in immune-related biological functions (Zhao et al., 2021). Thus, we have reason to believe that the regulatory mechanisms of small peptides are complex and diverse, and they may also respond to environmental changes and regulate growth and development as flexibly as protein-coding genes.

The MS-based peptide detection is a critical step for the identification of LSEPs. Given the incompatibility of ultracentrifuge filters with SDS-based protein extraction buffers, we tested two commonly used plant protein extraction buffers with modifications. The addition of HCl enhances protein extraction efficiency, consistent with results observed in animal cells (Ma et al., 2016). Buffer A demonstrates higher extraction efficiency for total protein extraction, while Buffer B performs better in extracting small proteins with molecular weights less than 20 kDa (**Fig. 2**). For MS data analysis, the sORF database yielded the highest number of identified peptides using PD search, showing a six-to-tenfold advantage over other methods. In summary, our study has successfully established a high-throughput and straightforward workflow for the identification of LSEPs. Additionally, it provides a valuable resource for future functional investigations of LSEPs in rice.

## Materials and Methods

### Plant materials and growth conditions

Rice (*Oryza sativa* ssp. *japonica*) Nipponbare were used in this study. Rice plants were grown in a growth chamber under the condition of 12-h light, 28°C, 70% relative humidity (RH), followed by 12-h dark, 23°C, 70% RH. Leaves of twenty-day-old rice were used for protein extraction. *Nicotiana benthamiana* plants were cultivated in a growth chamber under conditions of 16-h light/8-h dark, 22°C, 60% RH.

### RNA extraction, and lncRNA library construction

Total RNA was extracted using Trizol reagent kit (Invitrogen, Carlsbad, CA, USA) according to the manufacturer’s protocol. RNA quality was assessed on an Agilent 2100 Bioanalyzer (Agilent Technologies, Palo Alto, CA, USA) and checked using RNase free agarose gel electrophoresis. After total RNA was extracted, rRNAs were removed to retain mRNAs and ncRNAs. The enriched mRNAs and ncRNAs were fragmented into short fragments by using fragmentation buffer and reverse transcribed into cDNA with random primers. Second strand cDNA were synthesized by DNA polymerase I, RNase H, dNTP (dUTP instead of dTTP) and buffer. Next, the cDNA fragments were purified with QiaQuick PCR extraction kit (Qiagen, Venlo, The Netherlands), end repaired, poly (A) added, and ligated to Illumina sequencing adapters. Then UNG (Uracil-N-Glycosylase) was used to digest the second strand cDNA. The digested products were size selected by agarose gel electrophoresis, PCR amplified, and sequenced using Illumina HiSeqTM 4000 (or other platforms) by Gene Denovo Biotechnology Co. (Guangzhou, China).

### Ribo-seq library construction

The ribosomal profiling technique was carried out as reported previously(Zhu et al., 2023), with a few modifications as described below. The resuspended extracts in lysis buffer were transferred to new microtubes, pipetted several times and incubated on ice for 10 min. Then cells were triturated ten times through a 26-G needle.

The lysate was centrifuged at 20,000 g for 10 min at 4℃, and the supernatant was collected. To prepare RFs, 10 µL of RNase I (NEB, Ipswich, MA, USA) and 6 µL of DNase I (NEB, Ipswich, MA, USA) were added to 400 µL of lysate, which was then incubated for 45 min at room temperature with gentle mixing on a Nutator mixer. Nuclease digestion was stopped by adding 10 µL of SUPERase·In RNase inhibitor (Ambion, Austin, TX, USA). Size exclusion columns (illustra MicroSpin S-400 HR Columns; GE Healthcare; catalog no. 27- 5140-01) were equilibrated with 3 mL of polysome buffer by gravity flow and centrifuged at 600 g for 4 min at room temperature. 100 μL of digested RFs were added to the column and centrifuged at 600 g for 2 min. Next, 10 μL 10% (wt/vol) SDS was added to the elution, and RFs with a size greater than 17 nt was isolated according to the RNA Clean and Concentrator-25 kit (Zymo Research; R1017). rRNA was removed using the method reported previously(Guo et al., 2023). Briefly, short (50–80 bases) antisense DNA probes complementary to rRNA sequences was added to solution containing RFs, then RNase H (NEB, Ipswich, MA, USA) and DNase I (NEB, Ipswich, MA, USA) was added to digest rRNA and residual DNA probes. Finally, RFs were further purified using magnet beads (Vazyme, Nanjing, Jiangsu, China).

After obtaining ribosome footprints above, Ribo-seq libraries were constructed using NEBNext Multiple Small RNA Library Prep Set for Illumina (Catalog no. E7300S, E7300L). Briefly, adapters were added to both ends of RFs, followed by reverse transcription and PCR amplification. The 140-160 bp size PCR products were enriched to generate a cDNA library and sequenced using Illumina HiSeqTM X10 by Gene Denovo Biotechnology Co. (Guangzhou, China).

### Total and small peptide extraction

Same amount of rice leaves (1.5 g) were grounded into fine powder in liquid nitrogen, and resuspended with 6 ml pre-cooled extraction buffer containing 20 mM PMSF. HCl was added into the extracts with final concentration of 0 mM, 10 mM, 50 mM and 100 mM, and incubated on ice for 30 min with gently shaking. The samples were then boiled for 10 min, and mixed slowly on a shaker for 2 hours on ice. After centrifugation at 12,000 g for 30 min at 4°C, the supernatant was collected, and loaded on a 30-kDa MWCO filters (Millipore). After centrifugation at 15,000 g for 10 min, the remain samples (> 30 kDa) and flow through (< 30kDa) were all collected for further analysis.

### Mass spectrometry (MS) sample preparation

For sample preparation, we employed a filter-aided sample preparation (FASP) workflow, to remove the protein molecular weight higher than 30 kDa, as described before (Wiśniewski et al., 2009). The small peptides from FASP flow through were subsequently collected, cleaned, concentrated and enriched by a stop-and-go-extraction tips (StageTips) method (Rappsilber et al., 2007). The collected elute was completely dried in a speedvac centrifuge. Peptides were suspended in a buffer containing 2% acetonitrile and 0.1% TFA ready for MS analysis.

### LC-MS/MS and data analysis

The small peptides were analyzed by LC–tandem MS (LC–MS/MS) by combining an Easy-nLC1200 (Thermo Fisher Scientific) with Orbitrap Exploris 480 Mass Spectrometer (Thermo Fisher Scientific). A 100 μm × 2 cm trap column packed with Reprosil-Pur C18 5 μm particles (Dr Maisch GmbH) and a 75 μm × 25 cm analytical column packed with Reprosil-Pur C18 3 μm particles (Dr Maisch GmbH) were used to separate the peptides with mobile phase A (0.1% FA in water) and mobile phase B (0.1% FA in ACN) at a 60 min gradient: 6 to 23% B in 38 min, 23 to 32% B in 12 min, 32 to 80% B in 5 min, and then kept B at 80% for 5 min. The flow rate was set as 300 nl/min. The Orbitrap Exploris 480 Mass Spectrometer was operated in a data-dependent acquisition mode with a spray voltage of 2 kV and a heated capillary temperature of 320°C. MS1 data were collected at a high resolution of 60,000 with a mass range of 350 to 1800 m/z, the precursor intensity threshold was set at 5.0e^4^ and a maximum injection time of 20 ms. For each full MS scan, the top 10 most abundant precursor ions were selected for MS2 with an isolation window of 1.6 m/z and the higher energy collision dissociation with normalized collision energy of 30. MS2 spectrums were collected at a resolution of 15,000. The target value was 5e3 with a maximum fill time of 20 ms and a dynamic exclusion time of 30s.

Raw Data was processed and analyzed by MaxQuant (version 2.4.6.0) and PD 2.4 SP1 (Thermo Fisher Scientific, USA) with default settings. The in house generated searching database was used for searching as mentioned above. Human keratin was used as the contaminated database for proteomic searching. For small peptides database search, the digestion enzyme was set as unspecific. Oxidation (M) and Acetyl (Protein N-term) were specified as variable modifications. Q-value (FDR) cutoff on precursor and protein level was applied 1%. The processed data output were then exported to excel for further usage.

### sORFs prediction and custom database generation

The prediction of sORFs were carried out for each lncRNA using getorf from the EMBOSS package (Rice et al., 2000). The length of sORFs restricted a minimum length of 15 nt and a maximum length of 450 nt, and the alternative start codons were used, such as ATG, CTG, TTG. The physical position of sORFs on the chromosome was obtained using self-custom script. Duplicated sORFs were removed. Based on the nucleotide sequence of sORF, obtain its amino acid sequence and construct four custom peptide search libraries, including small peptide amino acid search library (sORF), only ribosomal binding small peptide amino acid search library (sORF_RF), small peptide amino acid and rice mRNA amino acid search library (sORF_mRNA), as well as ribosomal binding small peptide amino acid and mRNA amino acid search library (sORF_RF_mRNA).

### Vector construction and transient expression in *Nicotiana benthamiana*

Twelve putative LSEPs encoding lncRNAs with start coding of ATG, CTG and TTG were selected and individually fused to the N-terminus of the start codon-mutated *GFP* vectors (*pC3300-35S:ORF-mGFP-NOS*). The resulting constructs were transformed into Agrobacterium GV3101 competent cells. Tobacco (*N. benthamiana L.*) plants aged 4-5 weeks and grown in light were chosen as the host for infiltration. The *35S:p19* (a gene silencing suppressor) and *35S:ORF-mGFP* constructs were individually transformed into Agrobacterium tumefaciens cells (strain GV3101) by the freeze-thaw method, and *35S: mGFP* was used as a negative control, *35S:GFP* was used as a positive control. Transformed Agrobacterium cells were incubated at 28℃ for 2 days in LB medium containing 50 mg/ml kanamycin and 50 mg/ml rifampicin. Agrobacterium cells were resuspended in infiltration solution (10 mM MES, pH 5.7, 10 mM MgCl_2_, and 500 mM acetosyringone) and infiltrated into the abaxial side of tobacco leaves using a syringe as described previously. Infiltrated leaves were collected after 2 days for the sub-cellular localization assay or after 3-4 days for immuno-blot analysis.

### Rice protoplast transformation

Ten-day-old rice green seedlings were used for protoplast isolation. Stem and sheath tissues from 40-60 rice seedlings were cut into approximately 0.5 mm strips. The strips were immediately transferred into 0.6 M mannitol for a quick plasmolysis treatment, followed by enzymatic digestion in the dark with gentle shaking. The protoplasts were collected by filtration through 45 μm nylon meshes.

The CDSs of OsLSEPs or OsLSEPs without start condon (OsLSEPm) were fused with a GFP without start codon (mGFP), and under the control of ubiquitin promoter. The plasmids were transformed into rice protoplasts as described previously (Zhang et al., 2021). Fluorescence signals were detected by a Leica TCS SP8 confocal microscope at 24 hours post transformation.

### Immunoblot analysis

Total protein was extracted from leaf tissue after grinding in liquid nitrogen with protein extraction buffer (50 mM Tris-Cl pH 7.6, 5 mM EDTA, 5 mM EGTA, 2 mM DTT with protein phosphatase inhibitor and protease inhibitor cocktail). Protein accumulation was detected by anti-GFP antibodies (1:5000) (Abcam, Cambridge, UK). Signals were detected using a SuperSignal West Femto Trial Kit (Thermo Fisher Scientific), and visualized with a chemiluminescence detection system (Tianneng, Shanghai, China). Ponceau S staining of the membrane served as a loading control.

### Reads mapping and processing

All RNA-seq and Ribo-seq reads were processed and filtered using TrimGalore (Krueger, 2015) with default parameters and fastp (Chen et al., 2018), respecitvely. For Ribo-seq, reads with a length greater than 22 nt but less than 37 nt were retained, and paired-end reads were merged into single-end reads using fastp (Chen et al., 2018) with parameters “--length_required 23 --length_limit 36 --merge -overlap_len_require 23 --overlap_diff_limit 2”. The rRNA, tRNA and snoRNA sequences were downloaded from RNA central (https://rnacentral.org/search), and the retained Ribo-seq reads were aligned to these sequences using Bowtie2(Langmead and Salzberg, 2012) to get the unaligned reads. The unaligned reads were mapped to the rice genome (tigr7) using STAR(Dobin et al., 2013) with parameters “--outFilterMismatchNmax 2 --outFilterMultimapNmax 1 --alignEndsType EndToEnd” to get unique reads. The uniquely mapped reads were used for subsequent analysis. For RNA-seq, clean reads were mapped to rice genome (tigr7) using Hisat2 (Kim et al., 2015) with parameters “--rna-strandness RF”. The high quality (q30) uniquely mapped reads were used for subsequent analysis. FeatureCounts (Liao et al., 2014) was used to calculate the number of reads on features with default parameters. BEDTools (Quinlan and Hall, 2010) coverage was used to calculate the ribosomal footprint of predicted small open reading frames. The FPKM values were used to measure the expression level of the lncRNAs and mRNAs.

To evaluate the quality of Ribo-seq sequencing data, ribosome footprints (RF) length distribution, 3-nucleotide periodicity and metagene analysis were performed in self-custom script. For 3-nucleotide periodicity, the normalized reads coverage of the first base position at the upstream 40 bp and downstream 40 bp of transcript start and end sites for ribosomal footprints of the same length was calculated.

### Identification of rice lncRNAs

Strand-specific RNA-seq libraries were constructed using dUTP-based methods. The libraries were sequenced to get 150-bp paired-end reads on the illumina NovaSeq platform. A strict identification pipeline for lncRNA was used as previously described (Yuan et al., 2018). For each biological replicate, StringTie (Pertea et al., 2015) was used to de-novo assemble transcripts. All transcripts of the samples were merge and unified using StringTie merge (Pertea et al., 2015) and the annotation file of the rice reference genome (tigr7) was compared with the de-novo assemble transcripts using GffCompare (Pertea and Pertea, 2020). Only the transcript with the class code “i”, ”u”, “x”, “o” and “j” from GffCompare were retained. Transcripts with a length greater than 200bp were used for subsequent analysis. To evaluate the coding potential of transcripts, LGC (Wang et al., 2019), CPC2 (Kang et al., 2017) and pfam (Jones et al., 2014) with an E value cutoff of 1e-5 were used to predict non-coding transcripts. All the transcripts with coding-potential were removed and retained non-coding lncRNAs that appeared in all prediction tools.

### Bisulfite sequencing, DH-seq and ChIP-seq data processing

We downloaded previously published bisulfite sequencing (Zhang et al., 2018c), DH-seq (Zhang et al., 2012), ChIP-seq data for H3K36me3 (Zhang et al., 2012), H3K4me3 (He et al., 2010), H3K9ac (He et al., 2010), H3K27me3 (He et al., 2010), H3K9me2 (Zhao et al., 2020). The clean reads from bisulfite sequencing and ChIP-seq were mapped to tigr7 using Bismark (0.24.0) (Krueger and Andrews, 2011) and Bowtie2 (Langmead and Salzberg, 2012), respectively. The uniquely mapped reads were retained, and PCR duplicated reads were removed. For bisulfite sequencing, the extent of cytosine methylation was calculated with bismark_methylation_extractor in the Bismark program. For ChIP-seq, the MACS (2.2.8) (Zhang et al., 2008) program was used to identity read-enriched regions (peaks) with default parameters. Deeptools (3.5.1) (Ramírez et al., 2014) was used to calculate the signal intensity of markers in the upstream and downstream 2 kb regions of lncRNAs.

## Data availability

Data will be made available on request.

## Code availability

All R codes used for statistical analyses are available from the authors.

## Acknowledgements

This work was supported by the National Natural Science Foundation of China [grant numbers 32172420]; and The Fundamental Research Funds for the Central Universities, Grant/Award Number: [grant number KJJQ2024002 and KJJQ2024017].

## Author contributions

Y.Wang, J.Wu, and L.Wu conceived the project. H.Lin, L.Wei, S.Wang, T.Peng, J.Wu performed the experiments. Z.Chen performed bioinformatic analysis. W.Song performed the LC-MS/MS analysis. D.Wang, Y.Wu, L.Wei, J.Wu, and Y.Wang analyzed the data, Z.Chen, H.Lin, J.Wu, and Y.Wang worte the manuscript. All authors commented on the article.

## Supplementary information

**Supplemental Figure S1. The quality of sequencing data.**

(A) Total number of ribosome footprints falling near the beginning or end of CDSs. The 31-nucleotide RF reads and the combination across replicates are shown. The density of reads at each position was normalized. The x-axis represents the relative distance of each RF reads to the start codon or the stop codon. The deep blue, light green and light purple bars represent the first position of the RF reads mapped to 1st (expected), 2nd, 3rd reading frames, respectively.

(B) The three-nucleotide periodicity of 31-nucleotide RF reads. The frame percentage was inferred from the locations of the 5’ ends of the footprint at start and stop sites.

(C) The length distribution of ribosome footprint.

(D) The correlation heatmap of all samples. Colours range from blue to red representing correlation from high to low.

**Supplemental Figure S2. Characteristics of the start codon flanking sequence.** The probability of using four bases at each position of upstream 6 bp and downstream 5 bp of three start codons (AUG, CUG, UUG) in small peptides and mRNA (AUG).

**Supplemental Figure S3. Abundance characteristics of small peptides and mRNA in buffer A and buffer B for PD.**

(A-B) Abundance distribution of identified small peptides (A) and mRNA (B) in PD for different database. Green: total protein extracted using buffer A; purple: total protein extracted using buffer B.

(A) (C) The abundance of small peptides in different regions of the rice genome. Deep blue: total protein filtered using buffer A; light green: total protein filtered using buffer B. Wilcox-test was used for hypothesis testing. ns: p > 0.05; *: p < 0.05.

**Supplemental Figure S4. Characteristics of small peptides identified by MaxQuant.**

(A-B) Abundance distribution of identified small peptides (A) and mRNA (B) in different database by Maxquant. Green: total protein extracted using buffer A; purple: total protein extracted using buffer B.

(A) (C) Molecular weight of small peptides in different regions of the rice genome. Wilcox-test was used for hypothesis testing. *: p<0.05.

(B) (D) Length of translatable small open reading frames derived from different regions of the rice genome. Wilcox-test was used for hypothesis testing. *: p<0.05; **: p<0.01.

**Supplemental Figure S5. The association between lncRNA expression levels and epigenetic marks.**

(A) The percentage of lncRNAs associated with DH and histon modification in overall and translatable lncRNAs.

(B) Box plot of lncRNAs expression levels with or without DH and histion modification.

(C-D) Distribution of DH, histon modification, and DNA methylation over three types of lncRNAs, classified from high to low based on lncRNAs expression levels (FPKM >= 1). TSS, transcription start site; TES, transcription end site.

**Supplemental Table 1. Data quality control of LncRNA-seq and Ribo-seq.**

**Supplemental Table 2. Sequence of OsSLEPs and mutated OsSLEPs for validation.** Red colours indicated the predicted starting codon of OsSLEPs.

**Supplemental Data Set**

**Supplemental Data1.** Small peptides in the four databases identified by Proteome Discoverer.

**Supplemental Data2.** Abundance of small peptides in the four databases identified by MaxQuant.

**Supplemental Data3.** Peptides identified by MaxQuant in four databases.

**Supplemental Data4.** Expression level of LncRNAs and mRNAs.

**Supplemental Data5.** Identified lncRNAs regions and their classification.

**Supplemental Data6.** Molecular weight, length, isoelectric point and positional annotation of small peptides identified by Proteome Discoverer in the four databases.

**Supplemental Data7.** Molecular weight, length and positional annotation of the small peptides identified by MaxQuant in the four databases.

**Supplemental Data8.** Nucleotide and amino acid sequences of all identified small peptides.

**Supplemental Data9.** Annotations of all identified small peptides and expression levels of their corresponding lncRNAs.

**Supplemental Data10.** Types of epigenetic marks and expression levels of lncRNAs.

**Supplemental Data11.** Classification of lncRNAs based on multiple epigenetic marks and their corresponding expression levels.

**Supplemental Data12.** Classification of lncRNAs based on transposable element (TE).

**Supplemental Data13.** Ribosome binding levels of LncRNAs and mRNAs.

**Supplemental Data14.** Ribosome binding levels of all predicted sORFs.

