## Supplemental_Figures for "The multiomics landscape of small peptides encoded by long non-coding RNA-derived sORFs in rice"

### Supplement Figure1

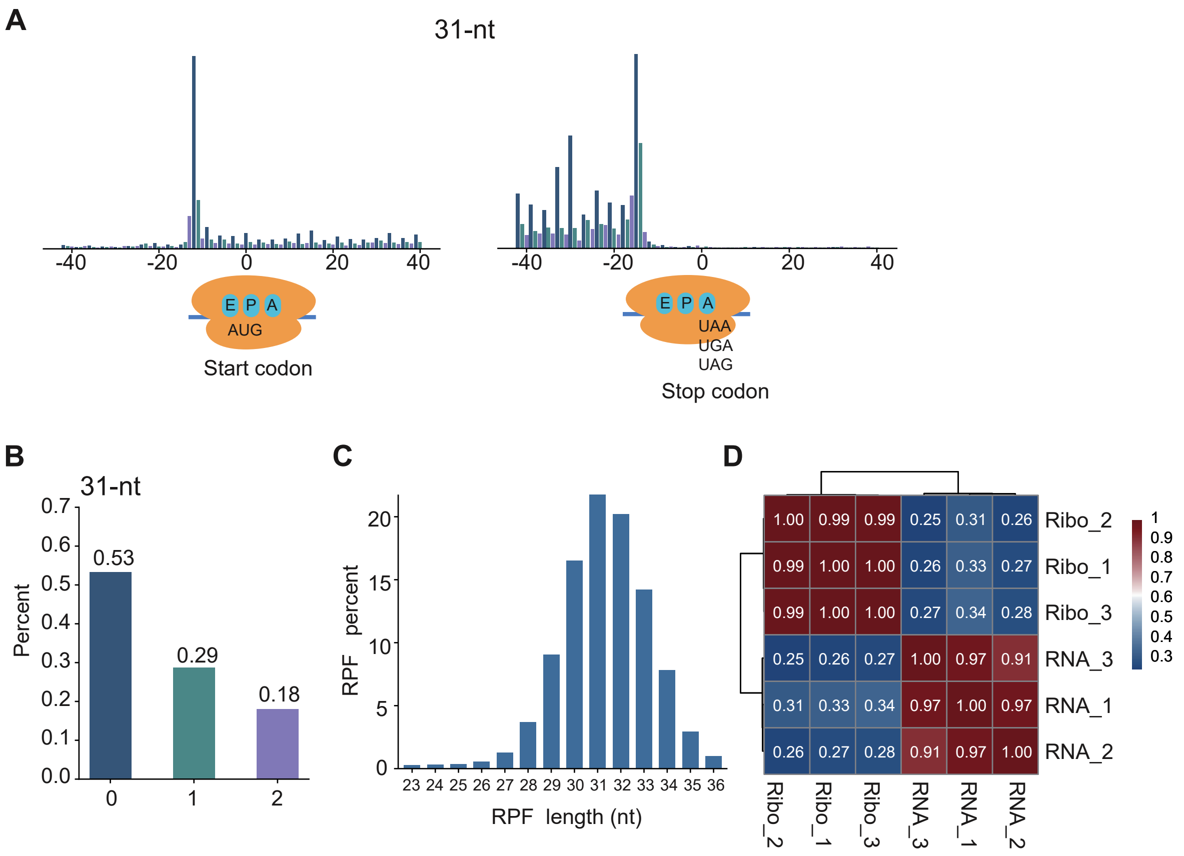


**Supplemental Figure S1. The quality of sequencing data.**

(A) Total number of ribosome footprints falling near the beginning or end of CDSs. The 31-nucleotide RF reads and the combination across replicates are shown. The density of reads at each position was normalized. The x-axis represents the relative distance of each RF reads to the start codon or the stop codon. The deep blue, light green and light purple bars represent the first position of the RF reads mapped to 1st (expected), 2nd, 3rd reading frames, respectively.

(B) The three-nucleotide periodicity of 31-nucleotide RF reads. The frame percentage was inferred from the locations of the 5' ends of the footprint at start and stop sites.

(C) The length distribution of ribosome footprint.

(D) The correlation heatmap of all samples. Colours range from blue to red representing correlation from high to low.

**Supplement Figure2**

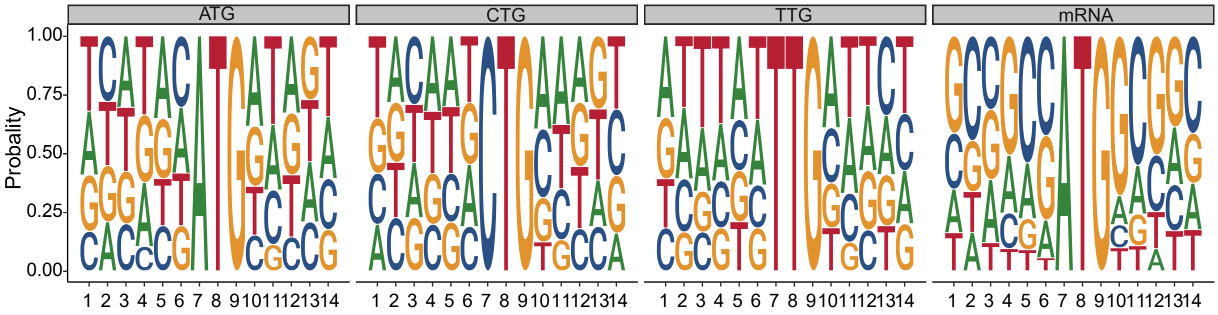


**Supplemental Figure S2. Characteristics of the start codon flanking sequence.**
The probability of using four bases at each position of upstream 6 bp and downstream 5 bp of three start codons (AUG, CUG, UUG) in small peptides and mRNA (AUG).

### Supplement Figure3

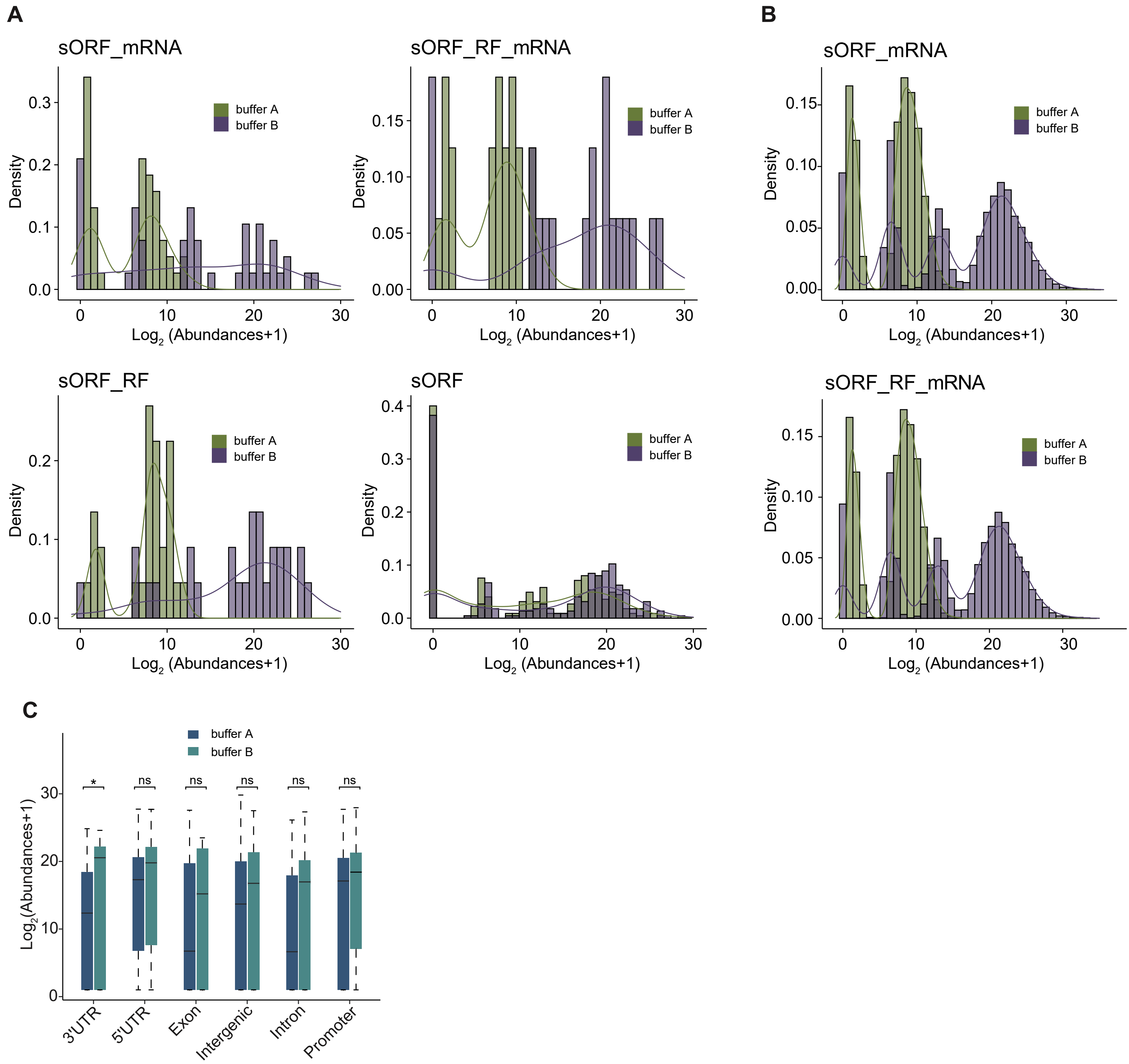


**Supplemental Figure S3. Abundance characteristics of small peptides and mRNA in buffer A and buffer B for PD.**

(A-B) Abundance distribution of identified small peptides (A) and mRNA (B) in PD for different database. Green: total protein extracted using buffer A; purple: total protein extracted using buffer B.

(C) The abundance of small peptides in different regions of the rice genome. Deep blue: total protein filtered using buffer A; light green: total protein filtered using buffer B. Wilcox-test was used for hypothesis testing. ns: $p > 0.05$; *: $p < 0.05$.

### Supplement Figure4

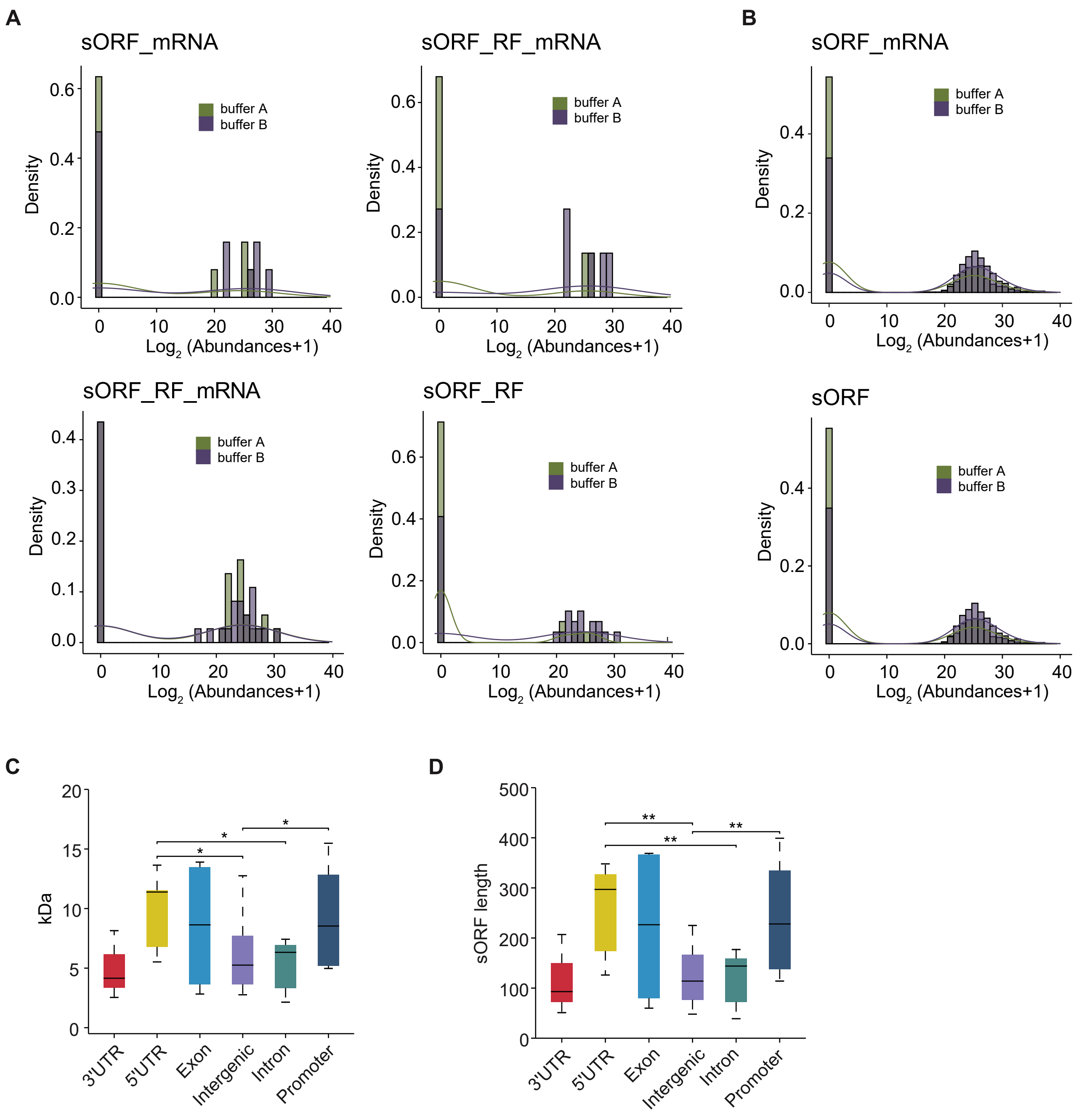


**Supplemental Figure S4. Characteristics of small peptides identified by MaxQuant.**
(A-B) Abundance distribution of identified small peptides (A) and mRNA (B) in different database by Maxquant. Green: total protein extracted using buffer A; purple: total protein extracted using buffer B.
(C) Molecular weight of small peptides in different regions of the rice genome. Wilcox-test was used for hypothesis testing. *: $p<0.05$.
(D) Length of translatable small open reading frames derived from different regions of the rice genome. Wilcox-test was used for hypothesis testing. *: $p<0.05$; **: $p<0.01$.

**Supplement Figure5**

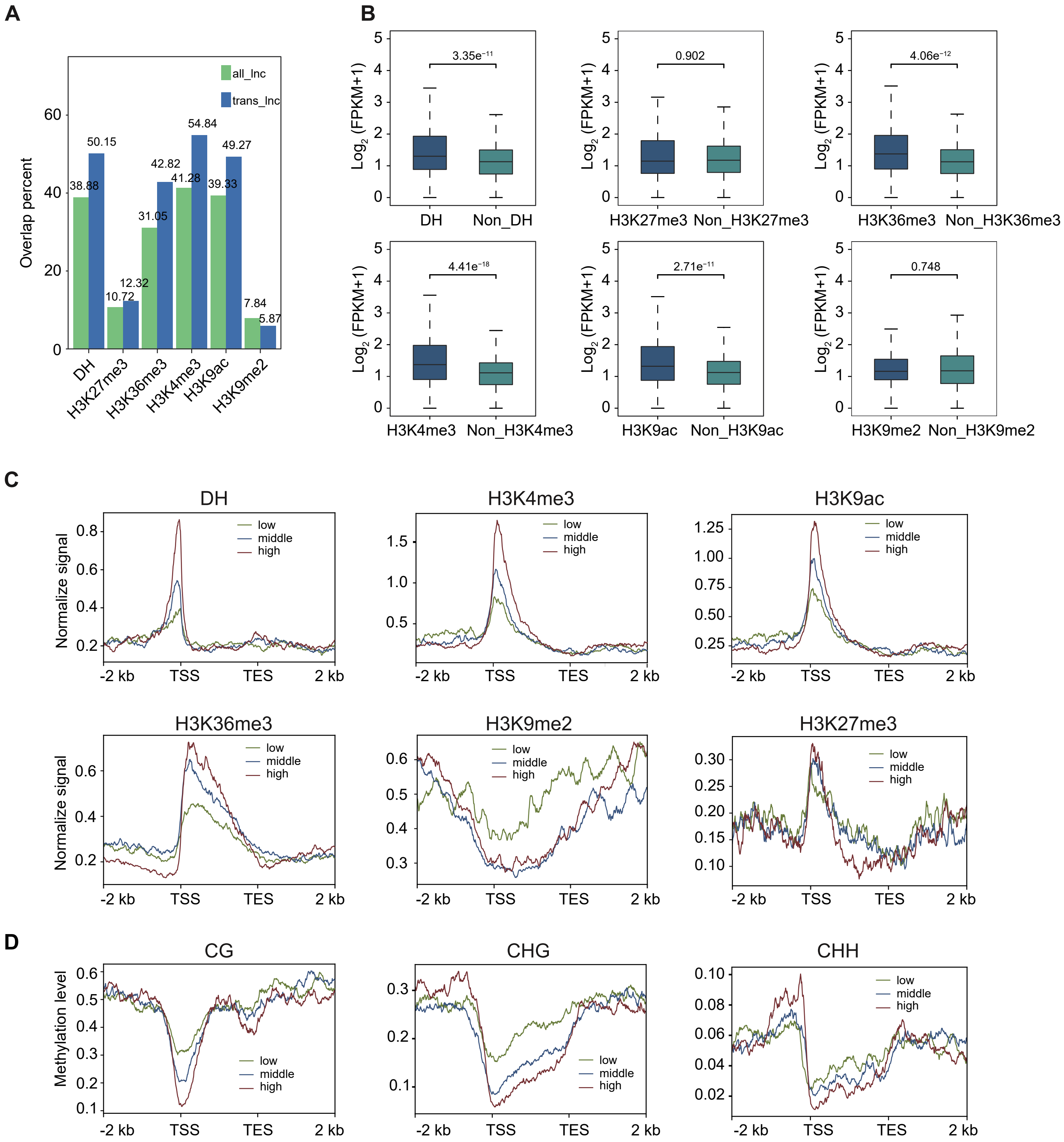


**Supplemental Figure S5. The association between lncRNA expression levels and epigenetic marks.**

(A) The percentage of lncRNAs associated with DH and histon modification in overall and translatable lncRNAs.

(B)Box plot of lncRNAs expression levels with or without DH and histion modification.

(C-D) Distribution of DH, histon modification, and DNA methylation over three types of lncRNAs, classified from high to low based on lncRNAs expression levels (FPKM >= 1). TSS, transcription start site; TES, transcription end site.
